# Brain-body timing gates conscious access across sensory systems

**DOI:** 10.64898/2026.05.08.723484

**Authors:** A. Wutz, R. Abt, F. Schmidt, T. Hartmann, M. Rogalla, M. Seifter, N. Weisz

## Abstract

Conscious awareness fluctuates with the brain’s internal state at the moment of perception, yet how these states are temporally coordinated remains unclear. Although such internal states are reflected in ongoing neural activity, their temporal structure may arise from intrinsic interactions between the brain and the body. To test this hypothesis, we simultaneously recorded neural, cardiac, and respiratory activity while participants detected auditory, tactile, or visual stimuli presented at perceptual threshold. Across sensory modalities, stimulus detection was preceded by transient bursts of alpha-band (8–13 Hz) activity localized to lateralized prefrontal and premotor regions. Crucially, these neural events were aligned with specific phases of the cardiac and respiratory cycles, revealing coordinated brain–body states at the time of perception. This cross-system coupling predicted whether sensory stimuli reached conscious awareness. Together, these findings show that brain–body synchrony acts as a temporal gate for conscious access across sensory systems.

## Introduction

Conscious access is governed by the observer’s internal state at the moment a stimulus occurs. What remains unclear is how these internal states are organized in time. Here, we test the hypothesis that brain state–dependent perception unfolds within a structured intrinsic temporal architecture, in which transient neural events—and their coordination with bodily rhythms—determine when external stimuli can enter awareness.

Decades of systems neuroscience have demonstrated that perception is not a passive reflection of sensory inputs. Instead, sensory signals are integrated within preconfigured neural networks whose activity is shaped by intrinsic circuitry and self-organized dynamics.^1–3^ Within this framework, brain states exhibit two defining features. First, they are not static but reverberate continuously over time.^4,5^ As large-scale network configurations evolve, so does perceptual experience, indicating that awareness is constructed internally rather than imposed by stimulation alone. Second, these states are not merely spontaneous but can be shaped by top-down control. Through regulation of the excitation–inhibition balance within cortical networks, the brain shapes its own readiness to process upcoming input, enabling observers to probabilistically bias internal states in anticipation of relevant events.^6–9^ A substantial body of work has established that such observer-dependent network dynamics predispose ongoing conscious experience, such that identical stimuli may be detected or missed depending on the brain’s momentary configuration.^10–12^

These findings marked a conceptual transition from stimulus-driven models of perception to state-dependent frameworks. Yet while the brain state-dependence of conscious access is now well established, it leaves open a fundamental question: are fluctuations in perceptual readiness simply stochastic variations in cortical excitability, or do they reflect an internally structured temporal architecture that coordinates when conscious access becomes possible?

Neural oscillations provide a candidate mechanism for organizing internal brain states in time. By routing activity through distributed neuronal assemblies and modulating synaptic interactions, oscillations support flexible transitions between network configurations while defining recurring temporal windows for perception and cognition.^13,14^ Because distinct neural populations generate oscillations at different frequencies, functional diversification may arise naturally in cortical networks. Among these rhythms, the brain’s dominant alpha frequency (8–13 Hz) has been closely linked to internally oriented brain states through its inhibitory influence on cortical processing.^15,16^ Alpha activity regulates cortical excitability to external stimuli and internally generated percepts,^17,18^ indexes internal shifts of selective spatial attention,^19,20^ and predicts whether an upcoming stimulus reaches awareness^21–24^ (for reviews see^25,26^). Together, these findings suggest that perceptual readiness does not fluctuate randomly but is shaped by intrinsic temporal structure, with alpha activity emerging as a central marker and mechanism of brain state–dependent perception.

If alpha activity reflects an intrinsic temporal architecture that gates perception, a critical question concerns how this architecture is instantiated in real time. Most evidence relies on trial-averaged measures of oscillatory power, which may obscure the fine-grained temporal organization of neural events at the single-trial level. Recent advances suggests that oscillatory activity often manifests as transient, burst-like events rather than sustained rhythms.^27–30^ Trial-averaged power may therefore conflate the rate, duration or timing of temporally resolved inhibitory events with sustained amplitude changes. The precise timing of such bursts may be more informative than their overall magnitude, as they may define momentary windows of elevated cortical excitability and perceptual readiness.^31,32^ Identifying when alpha bursts occur therefore provides a mechanistic means of testing whether structured neural timing of intrinsic neural events, rather than slower stochastic fluctuations of global oscillatory power, governs conscious access at specific moments in time.

Characterizing alpha activity as temporally resolved bursts reframes perceptual gating as a problem of timing. The critical issue then becomes what coordinates the occurrence of these intrinsic events. An increasingly influential perspective highlights the central role of brain–body coupling in organizing neural timing.^33–35^ Cardiac and respiratory rhythms impose intrinsic temporal structure on neural activity and have been shown to interact with perceptual sensitivity and decision-making within individual sensory modalities.^36–41^ These bodily rhythms generate continuous fluctuations in afferent signaling and arousal that may act as a global timing signal coordinating distributed neural processes. Moreover, brain–body coupling seems conserved across species,^42,43^ suggesting an evolutionarily advantageous mechanism for aligning perception and action in time. Rather than reflecting local neural variability alone, the timing of inhibitory bursts may therefore be scaffolded by organism-wide physiological dynamics. Such brain-body coordination may in turn gate conscious access.

The aim of this study was to determine whether perceptual detection across sensory modalities is predicted by the timing of intrinsic alpha-band dynamics and their coordination with cardiac and respiratory rhythms. To address this question, we investigated auditory, tactile, and visual perception within the same participants and experimental framework. Brief stimuli (50 ms) were presented at individually titrated perceptual thresholds to achieve comparable proportions of detected and missed trials, thereby dissociating perceptual outcome from stimulus intensity (Fig. 1A). This multimodal design enabled us to distinguish supramodal mechanisms of perceptual readiness that generalize across sensory systems from modality-specific sensory processing.^44^ Concurrent recordings of magnetoencephalography (MEG), electrocardiography (ECG), and respiratory activity allowed precise characterization of neural and bodily dynamics at the time of perception.

**Figure 1.**
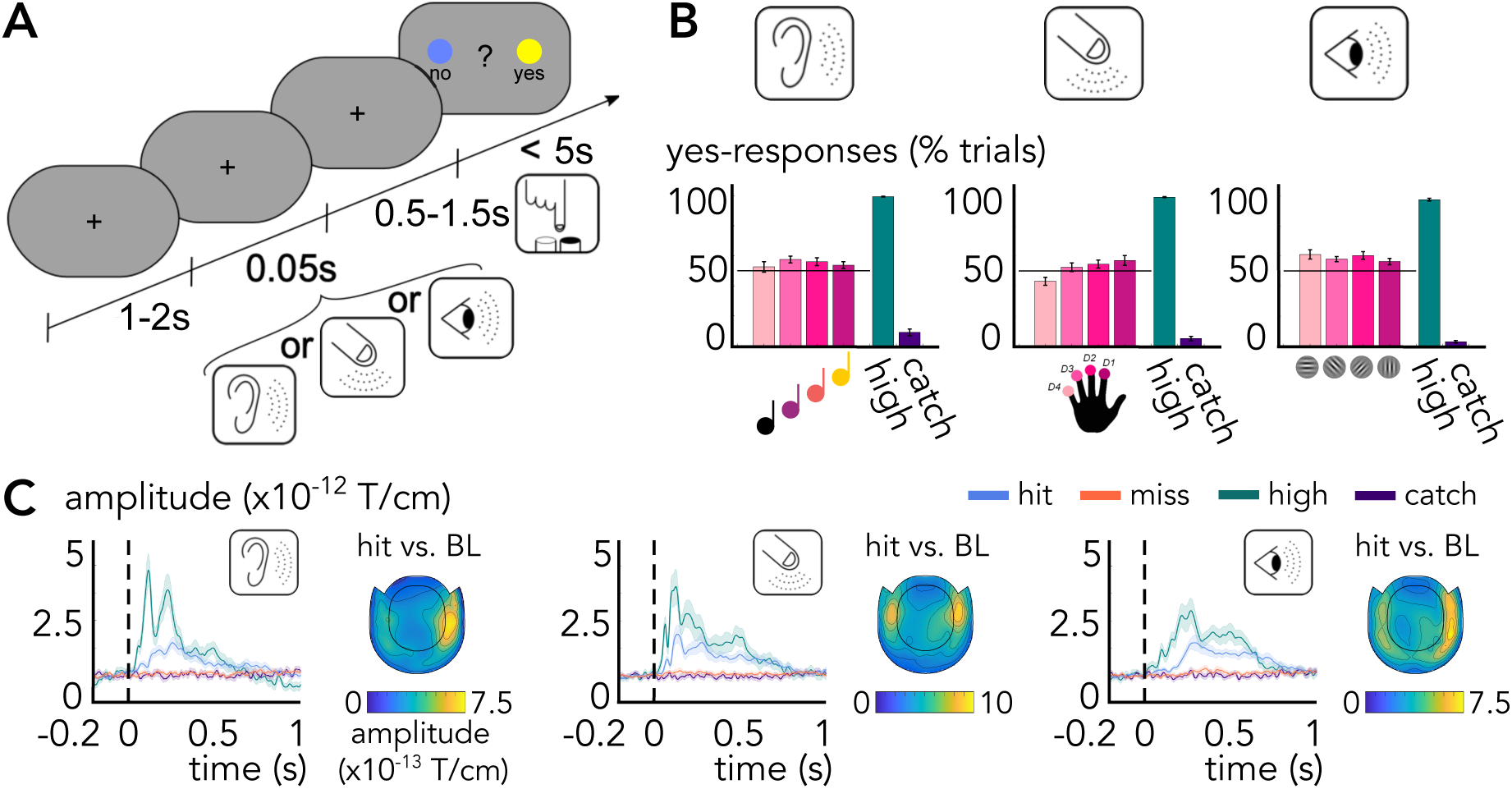
Task sequence, behavioral data and stimulus-evoked MEG fields **A** Task sequence of the multimodal, near-threshold experiment. All auditory, tactile and visual stimuli were presented for 50 ms in left external space, i.e. in the left ear, hand or visual hemifield. The stimulus intensities were titrated to reach approximately equal proportions of perceived vs. not perceived reports in a yes-no detection task. **B** Behavioral performance in percent yes-reports for near-threshold stimuli (pink hues), high signal- (green) and no signal- trials (purple). It was close to the intended 50% yes-reports (see horizontal black line) for all near-threshold stimuli. Error bars represent standard errors of the mean (SEM) for repeated-measures designs. **C** MEG activity time-locked to stimulus onset per modality for detected (blue), missed (red), high signal- (green) and no signal-trials (purple). Shaded error regions represent SEM for repeated-measures designs. The MEG time series were averaged across the sensors with the greatest evoked activity per modality, as shown by the inset topographies.

We first assessed behavioral performance and stimulus-evoked neural responses as confirmatory benchmarks, and replicated established pre- and post-stimulus alpha power effects to anchor our findings within the literature on state-dependent perception. Building on this foundation, we then examined single-trial alpha burst timing to test whether transient intrinsic events predict perceptual outcome across modalities. Finally, we investigated whether alpha burst occurrence was coordinated with cardiac and respiratory rhythms, and whether such brain–body coupling enhanced prediction of conscious detection. This analytical progression enabled us to examine whether conscious access across sensory systems is structured not only by intrinsic neural dynamics but also by coordinated brain–body rhythms.

## Results

### Behavioral data and stimulus-evoked MEG activity followed experimental expectations

Behavioral performance closely followed the intended experimental design. On near-threshold trials, detection rates were close to the targeted 50% trials across all three sensory modalities (mean yes-responses ± SD, % of trials: auditory, 55 ± 9%; tactile, 52 ± 8%; visual, 59 ± 9%). In contrast, detection on high-signal trials with clearly suprathreshold stimuli was near ceiling (auditory, 99 ± 1%; tactile, 99 ± 2%; visual, 98 ± 4%), while false-alarm rates on catch trials, in which no stimulus was presented, were low (auditory, 9 ± 10%; tactile, 5 ± 6%; visual, 3 ± 4%; Fig. 1B). Signal detection analyses confirmed high perceptual sensitivity for individually titrated near-threshold stimuli (mean d′ ± SD: auditory, 1.7 ± 0.4; tactile, 1.9 ± 0.5; visual, 2.2 ± 0.5), accompanied by relatively conservative response criteria (mean c ± SD: auditory, 0.7 ± 0.4; tactile, 0.9 ± 0.3; visual, 0.9 ± 0.2). This behavioral pattern was mirrored in stimulus-evoked MEG responses. Clear post-stimulus evoked fields were observed for detected and high-signal trials, whereas miss and catch trials elicited minimal evoked activity (Fig. 1C). Consistent with established sensory processing pathways, evoked responses were strongest over modality-specific sensory regions and lateralized mostly contralaterally to stimulation, which was in left external space, i.e. in the left ear, hand or visual hemifield.

### Alpha power distinguishes perceptual outcomes and shows cross-modal consistency

We then analyzed pre-stimulus alpha power and post-stimulus alpha desynchronization to replicate well-established markers of brain state–dependent perception, providing a reference point against which to interpret subsequent analyses. Time–frequency resolved MEG analyses revealed robust power fluctuations with a similar temporal structure across modalities that were maximal in the alpha-frequency range (8–13 Hz), consistent with our a priori hypothesis and motivating the focus on alpha in subsequent analyses (Fig. 2A). These effects formed a structured temporal pattern spanning pre- and post-stimulus periods. In the pre-stimulus interval, overall alpha power increased several hundred milliseconds before stimulus onset, predominantly over bilateral frontal sensors. When comparing perceptual outcomes, pre-stimulus alpha power did not significantly differ between detected and missed trials in the auditory modality and showed only a trend-level effect for touch (p < .08). In contrast, and consistent with previous work,^21,22^ pre-stimulus alpha power was significantly higher before missed compared with detected visual stimuli (p < .002; Fig. 2B). Post-stimulus intervals revealed a complementary and highly consistent pattern across modalities. Detected stimuli were followed by pronounced alpha (and beta) desynchronization, reflecting the canonical event-related power decrease associated with sensory-motor processing and cortical engagement.^15,45,46^ For alpha, this desynchronization was markedly attenuated or absent for missed stimuli, resulting in significant post-stimulus alpha power differences that were spatially specific and predominantly ipsilateral to the stimulated side (auditory, *p* < .01; tactile, *p* < .002; visual, *p* < .002; Fig. 2B). Together, this characteristic time–frequency profile—sustained pre-stimulus alpha activity followed by robust post-stimulus alpha desynchronization—indicates a high degree of cross-modal generality in alpha-band dynamics associated with perceptual detection across audition, touch, and vision.

**Figure 2.**
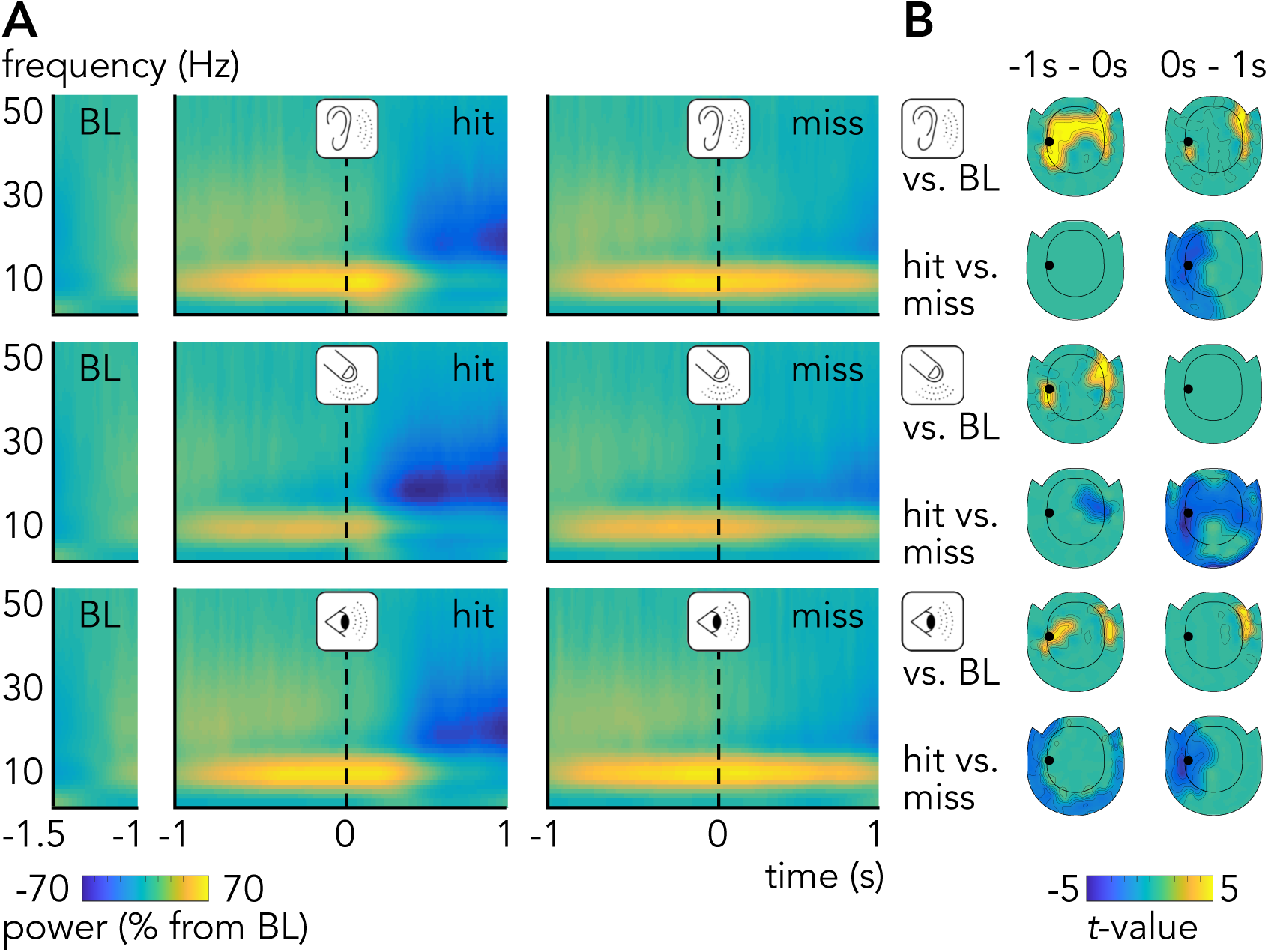
Different alpha power per perceptual outcome but similar power across the senses **A** Time-frequency maps for oscillation power vs. pre-trial baseline (-1.5s to -1s) for detected (left) and missed (right) auditory (upper row), tactile (middle row) and visual stimuli (lower row). The time-frequency maps were plotted for one representative MEG gradiometer sensor (black dot on sensor-level topographies in **B**). **B** Sensor-level topographies for alpha-band power (8-13 Hz) in pre- (left) and post-stimulus intervals (right) per modality. The upper rows show repeated-measures *t*-statistics between all near-threshold trials per modality vs. pre-trial baseline (Bonferroni-corrected). The lower rows show repeated-measures *t*-statistics between hit- vs. miss-trials per modality (cluster-corrected). All topographies are masked at a corrected threshold of *p* < .05.

### Pre-stimulus alpha burst timing predicts perception across sensory systems

While post-stimulus alpha (and beta) desynchronization has been linked to sensory and motor response processes,^15,45,46^ pre-stimulus alpha activity is thought to reflect ongoing internal states that bias subsequent perceptual outcomes.^15–26^ To characterize the fine-grained temporal structure underlying these effects, we performed a single-trial oscillatory burst analysis. This approach identifies transient increases in oscillatory activity on individual trials with high temporal precision.^27–30^ Burst analyses quantify neural activity relative to a local, trial-specific baseline, effectively capturing the prominence of brief increases above the surrounding activity level rather than their absolute amplitude. A sensitivity analysis using five independent burst detection algorithms yielded qualitatively and quantitatively consistent results, indicating that the observed effects are robust to alternative detection methods and not driven by baseline differences in overall power between conditions (see Methods, Fig. 3A and Supplementary Fig. 1).

**Figure 3.**
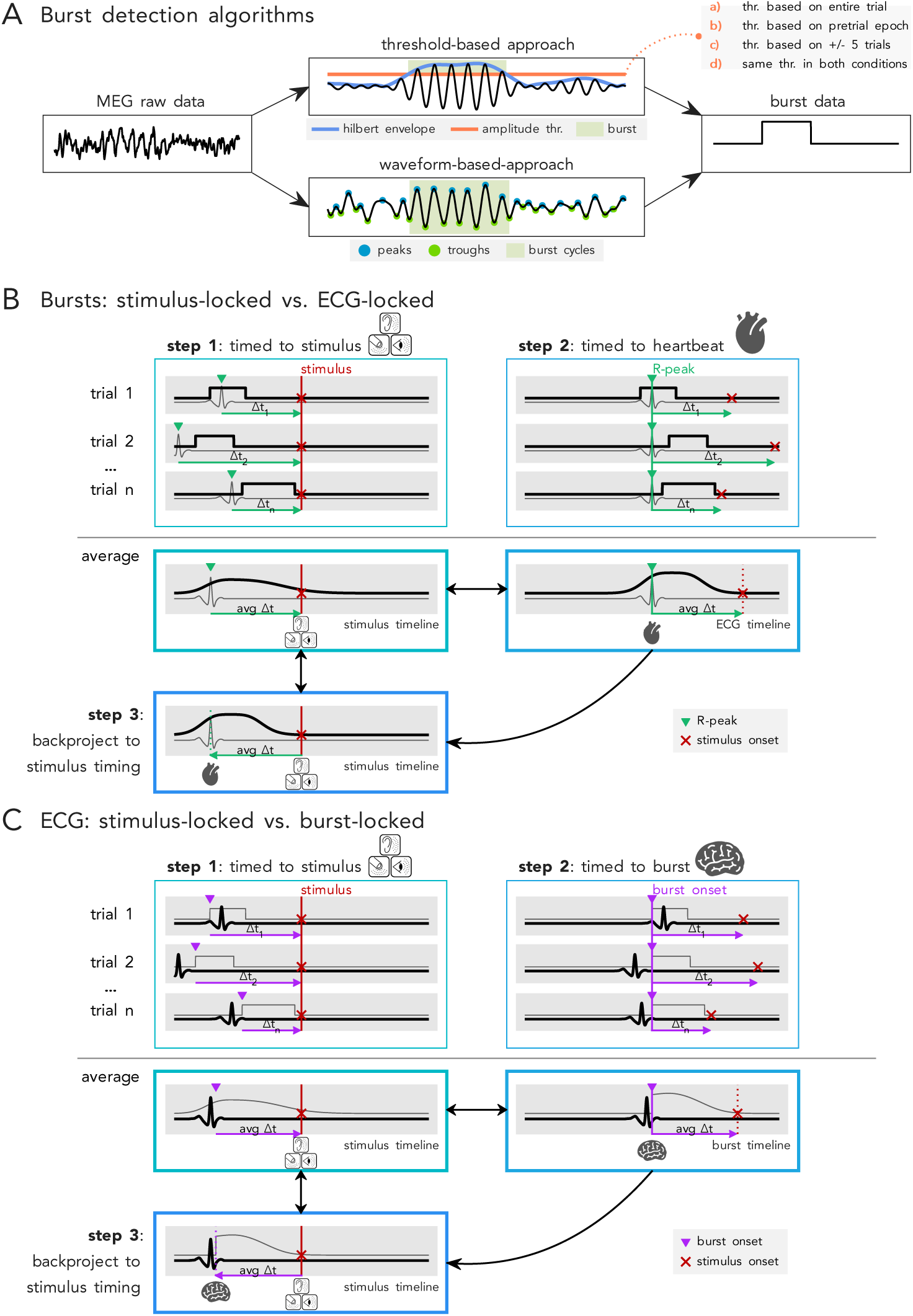
Analytical framework for burst detection and neural-physiological temporal alignment **A** Schematic depiction of burst detection algorithms. Four different threshold-based (a-d) and one waveform-based method were evaluated. **B** Forward logic: temporal alignment of burst activity to heartbeats. First, neural burst time series are aligned to stimulus onset (step 1). Then, neural burst time series are expressed relative to cardiac cycles by aligning trials to R-peak times and averaging across trials (step 2). Finally, the resulting average burst rate is backprojected into stimulus time based on the average pre-stimulus cardiac event timing (step 3). **C** Reverse logic: temporal alignment of cardiac signals to burst onsets. First, cardiac time series are aligned to stimulus onset (step 1). Then, cardiac time series are expressed relative to neural activity by aligning trials to burst onset times and averaging across trials (step 2). Finally, the resulting average cardiac signals are backprojected into stimulus time based on the average pre-stimulus burst onset timing (step 3). For clarity, only cardiac signals are shown in **B** and **C**. The procedure for respiratory signals is analogous using the respiration peaks, instead of the R-peaks.

We computed burst rates as the percentage of trials in which a burst is present within a given time window. Across modalities, alpha bursts occurred on a substantial proportion of trials within the -1 to 1 s interval centered on stimulus onset (mean ± SD: auditory, 56 ± 18%; tactile, 45 ± 20%; visual, 55 ± 18%). Bursts consistently began in the pre-stimulus interval (mean onset ± SD: auditory, −180 ± 80 ms; tactile, −182 ± 91 ms; visual, −195 ± 64 ms), lasted approximately five alpha cycles (mean duration ± SD: auditory, 520 ± 78 ms; tactile, 493 ± 81 ms; visual, 543 ± 98 ms), and often extended into the post-stimulus period (mean offset ± SD: auditory, +340 ± 132 ms; tactile, +311 ± 117 ms; visual, +348 ± 113 ms; Fig. 4). Consistent with power-based analyses, alpha burst rates were highest over bilateral frontal sensors. Post-stimulus burst rates were significantly higher for missed than for detected stimuli across all modalities (all p < .002), mirroring post-stimulus alpha desynchronization effects. Crucially, the burst analysis also revealed complementary pre-stimulus effects at similar, predominantly ipsilateral sensors. Alpha burst rates were significantly higher before detected than missed stimuli for audition (p < .002), touch (p < .01), and vision (p < .04; Fig. 4). Single-trial analyses further showed that, for detected stimuli, alpha burst onsets occurred earlier and were more temporally clustered in the pre-stimulus interval than for missed stimuli across all modalities (all p < .004). Importantly, burst durations were comparable between detected and missed trials, indicating that condition differences were not primarily driven by changes in event duration, but instead reflected differences in burst timing and its temporal variability (see Supplementary Fig. 1). These findings indicate that alpha burst timing distinguishes perceptual outcomes already before stimulus onset across sensory systems.

**Figure 4.**
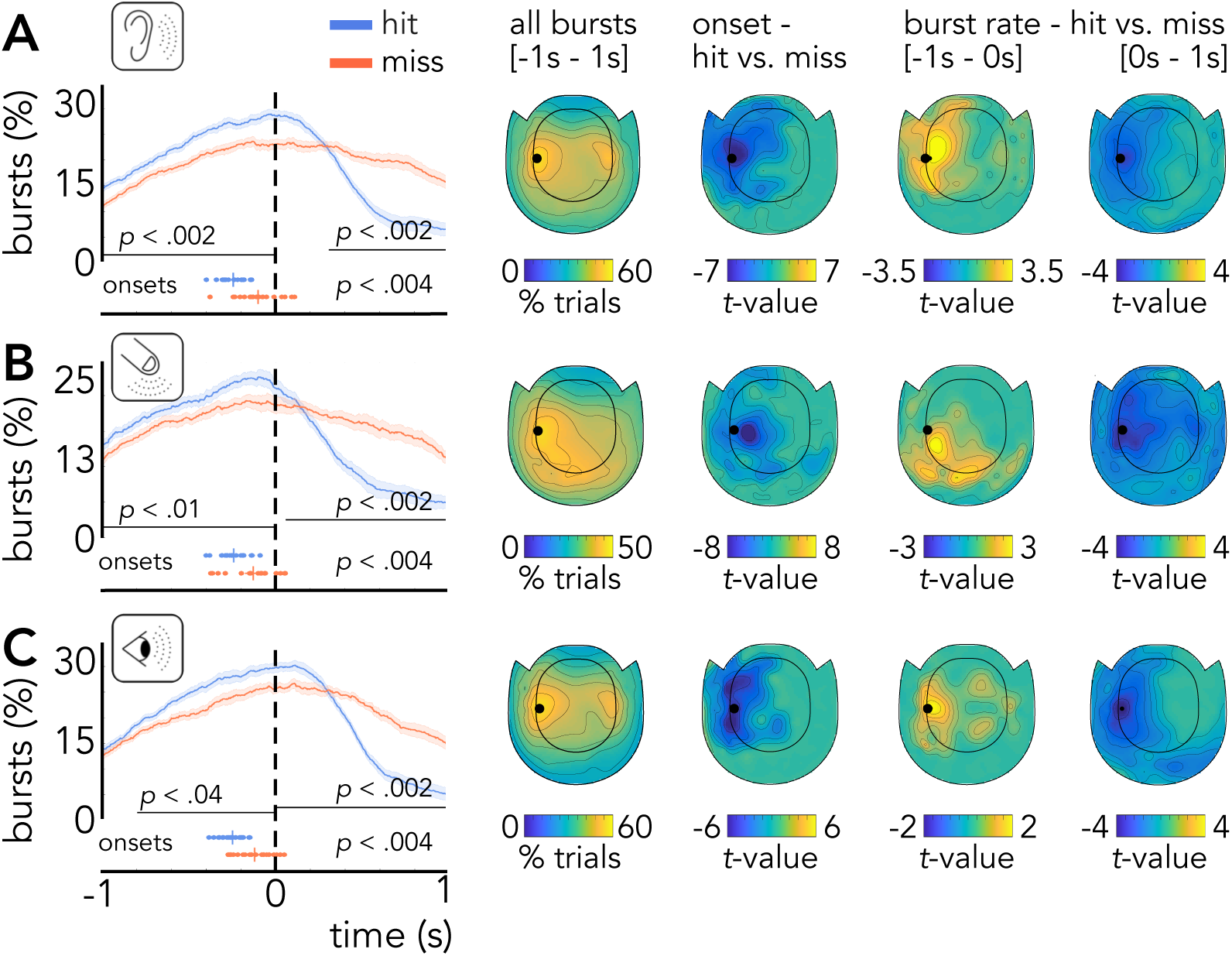
Different alpha burst activity per perceptual outcome but similar burst activity across the senses **A** Burst rates in % trials over time in the alpha band (8-13 Hz) for detected (blue) and missed (red) auditory stimuli on a representative MEG gradiometer sensor (black dot on sensor-level topographies on the right). Shaded error regions represent SEM for repeated-measures designs. Inset circles show the average burst onsets per participant and perceptual outcome. Inset vertical lines show the mean onsets. *P*-values denote significance levels for repeated-measures *t*-statistics (cluster-corrected). The sensor-level topographies on the right show all burst occurrences in the interval between -1s to 1s (in percent trials), repeated-measures *t*-statistics between hit- vs. miss-trials for the burst onsets, as well as for the pre-stimulus and post-stimulus burst rates (all cluster-corrected). The topographies showing statistics are masked at a corrected threshold of *p* < .05. **B** and **C** as in **A** but for tactile or visual stimuli.

### Pre-stimulus alpha burst timing is coordinated by cardiac and respiratory cycles

Building on this supramodal consistency, we pooled trials across modalities to examine whether the predictive timing of pre-stimulus alpha bursts is coordinated by intrinsic brain–body rhythms. To test this, we compared stimulus-timed analyses with heart- and respiration-timed analyses. Stimulus-timed analyses replicated the modality-specific results reported above: alpha burst rates were higher for detected than missed stimuli before stimulus onset (*p* < .006), and higher for missed than detected stimuli after stimulus onset (*p* < .002; Fig. 5A). These effects were driven by earlier burst onsets for detected stimuli over ipsilateral sensors (*p* < .002). To examine neural activity relative to internal physiological timing, we re-expressed burst time series in a cardiac- or respiration-centered reference frame. Specifically, for each trial, the time axis was temporarily aligned to the last heartbeat (R-peak) or respiration peak preceding stimulus onset, and burst dynamics were computed in this interoceptive time frame. The resulting time series were then projected back into stimulus time using each participant’s mean cardiac or respiratory timing before stimulus onset (see Methods and Fig. 3B). This analysis strategy enables direct comparison between heart- and respiration-timed data with stimulus-locked analyses, while preserving the original temporal relationship to stimulus onset (Fig. 5A–C). Potential contributions of cardiac field artifacts were mitigated using multiple complementary preprocessing and validation approaches. Control analyses confirmed progressive reduction of cardiac-related signal contributions without compromising the underlying MEG data (see Methods and Supplementary Fig. 2). On average, the last heartbeat occurred at −436 ± 41 ms and the last respiration peak at −473 ± 88 ms before stimulus onset, with no significant timing differences between detected and missed trials (heart: *p* < .07; respiration: *p* < .38). This suggests that differences in perceptual outcome are not driven by cardiac or respiratory timing alone, but may instead depend on their alignment with ongoing neural dynamics, which we examine in the following analysis. Neural burst activity increased time-locked to both cardiac and respiratory peaks and differed by perceptual outcome, as reflected in burst rates (heart: pre-stimulus *p* < .002, post-stimulus *p* < .002; respiration: pre-stimulus *p* < .008, post-stimulus *p* < .002) and burst onset times (heart and respiration: both *p* < .002). Relative to cardiac and respiratory cycles, alpha bursts occurred earlier and with reduced temporal variability on detected compared with missed trials. On detected trials, burst activity largely terminated before stimulus onset, whereas on missed trials it extended into post-stimulus periods. The shared ipsilateral topographies across stimulus-, heart-, and respiration-timed analyses suggest common neural generators (Fig. 5D–F). Source reconstruction confirmed this convergence (Fig. 5G). Consistent with sensor-level effects, overall burst activity was strongest in bilateral premotor and supplementary motor regions (Brodmann Area (BA) 6), with additional contributions from visual cortex (BAs 19 and 37). The ipsilateral differences between detected and missed trials found on the sensor-level localized to left supramarginal gyrus (BA 40), premotor/supplementary motor cortex (BA 6), superior temporal gyrus (BA 22), dorsolateral prefrontal cortex (BAs 9 and 46), and inferior frontal gyrus (BAs 45 and 47), evident in burst onset timing (*p* < .002), pre-stimulus burst rates (*p* < .016 and *p* < .044), and post-stimulus burst rates (*p* < .002). These regions are implicated in top-down attentional control and perceptual readiness, consistent with a supramodal mechanism.^47,48^ Contralateral effects were observed in visual cortex (BAs 18 and 19), superior parietal cortex (BA 7), premotor cortex (BA 6), temporal cortex (BA 38), and prefrontal regions (BAs 8, 9, and 10). Importantly, power increases following heartbeats - extending into the alpha-to-low-beta range - were also present during rest (Supplementary Fig. 3), indicating that brain–cardiac coupling reflects an intrinsic process rather than a task-specific effect. Convergent task and rest effects localized to anterior cingulate cortex (BA 32), frontal eye fields (BA 8), and dorsolateral prefrontal cortex (BAs 9 and 46), regions associated with interoception and self-monitoring.^49,50^

**Figure 5.**
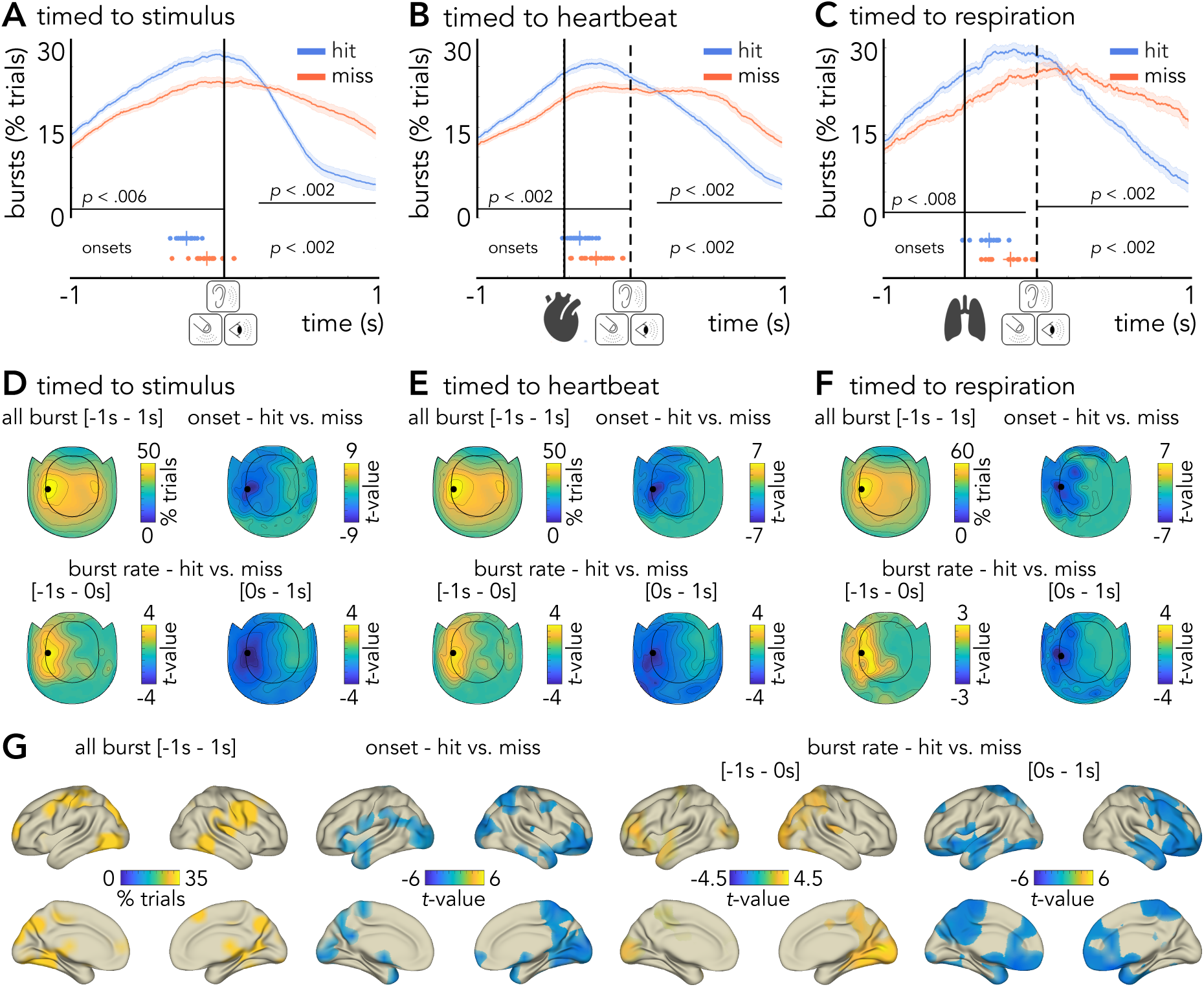
Burst rates per perceptual outcome timed to stimuli, heartbeats and breathing **A** Burst rates in % trials over time in the alpha band (8-13 Hz) for detected (blue) and missed (red) trials on a representative MEG gradiometer sensor (black dot on sensor-level topographies in **D-F**) time-locked to stimulus onset (straight vertical line). Shaded error regions represent SEM for repeated-measures designs. Inset circles show the average burst onsets per participant and perceptual outcome. Inset vertical lines show the mean onsets. *P*-values denote significance levels for repeated-measures *t*-statistics (cluster-corrected). **B** and **C** as in **A** but time-locked to R-peaks or respiration peaks, respectively, and then shifted to the average peak times per participant (straight vertical lines) before stimulus onset (dotted vertical lines). See Figure 3B for a schematic overview of the method. **D** Sensor-level topographies for all burst occurrences in the interval between -1s to 1s (in percent trials), for repeated-measures *t*-statistics between hit- vs. miss-trials for the burst onsets, as well as for the pre-stimulus and post-stimulus burst rates (all cluster-corrected). The topographies showing statistics are masked at a corrected threshold of *p* < .05. **E** and **F** as in **D** but for the ECG- or respiration-timed burst activity. **G** Source-level distributions for the stimulus-locked burst occurrences in the interval between -1s to 1s (in percent trials and masked at the upper 10%-voxels), for repeated-measures *t*-statistics between hit- vs. miss-trials for the burst onsets, as well as for the pre-stimulus and post-stimulus burst rates (all cluster-corrected). The source reconstructions showing statistics are masked at a corrected threshold of *p* < .05.

### Cardiac and respiratory rhythms align with pre-stimulus alpha burst timing

Finally, we tested whether cardiac and respiratory activity was more strongly aligned to intrinsic neural events than to external stimulus timing. To do so, we applied the same analytical framework as above but reversed the reference frame: instead of aligning neural activity to cardiac or respiratory events, we realigned ECG and respiration signals to alpha burst onsets on a trial-by-trial basis. The resulting physiological time series were then projected back into stimulus time using each participant’s mean burst onset timing separately for detected and missed trials, enabling direct comparison with stimulus-locked analyses while preserving the original temporal relationship to stimulus onset (see Methods and Fig. 3C). Stimulus-timed ECG showed no consistent modulation across systole and diastole, whereas burst-aligned ECG revealed a clear biphasic pattern (Fig. 6A,B). Alpha burst onsets occurred at a consistent cardiac phase, exhibiting strong ECG phase coherence across trials (permutation test, *p* < .001) and participants (Rayleigh test, *z* = 11, *p* < 4 × 10^-6^), an effect absent at stimulus onset (Fig. 6C). Burst onsets occurred at similar cardiac phases for detected and missed trials; however, because bursts occurred earlier in absolute time on detected trials, the cardiac phase relative to stimulus onset differed significantly between perceptual outcomes (mean phase difference 41°, 95% CI [10°, 73°]; m-test vs. 0°: *p* < .015; Rayleigh test for coherence: z= 6.3, *p* < .002; Fig. 6D). On detected trials, cardiac activity was already approaching diastole at stimulus onset, whereas on missed trials it coincided with peak systole. This phase relationship was absent in stimulus-timed analyses and confirms strong coupling between neural and cardiac activity. Parallel analyses of resting-state data revealed comparable coupling, further supporting its intrinsic nature (Supplementary Fig. 4). Burst onsets in both task- and rest data exhibited a consistent phase preference within the cardiac cycle but were significantly offset from the R-peak (m-test for the phase difference vs. 0° for task: *p* < .0005; for rest: *p* < .038). Given the intrinsic coupling between cardiac and respiratory rhythms^51^ (see also Supplementary Fig. 5), we examined respiration as a potential mediator. Participants tended to suppress breathing out during pre-stimulus periods, with no significant stimulus-timed differences between detected and missed trials (Fig. 6E). In contrast, burst-aligned respiratory signals revealed significant timing differences (*p* < .004; Fig. 6F): on detected trials, exhalation began earlier and coincided with burst onset, such that participants were already in the exhalation phase at stimulus onset, whereas exhalation occurred later on missed trials. Notably, this difference reflects an earlier shift in exhalation onset relative to burst timing, such that condition differences become most apparent during and after stimulus presentation despite arising in pre-stimulus intervals. In sum, these findings reveal intrinsic temporal coupling between neural alpha bursts, cardiac activity, and respiration. This neuro–cardio–respiratory coordination, largely invisible in traditional stimulus-locked analyses, robustly predicted perceptual detection.

**Figure 6.**
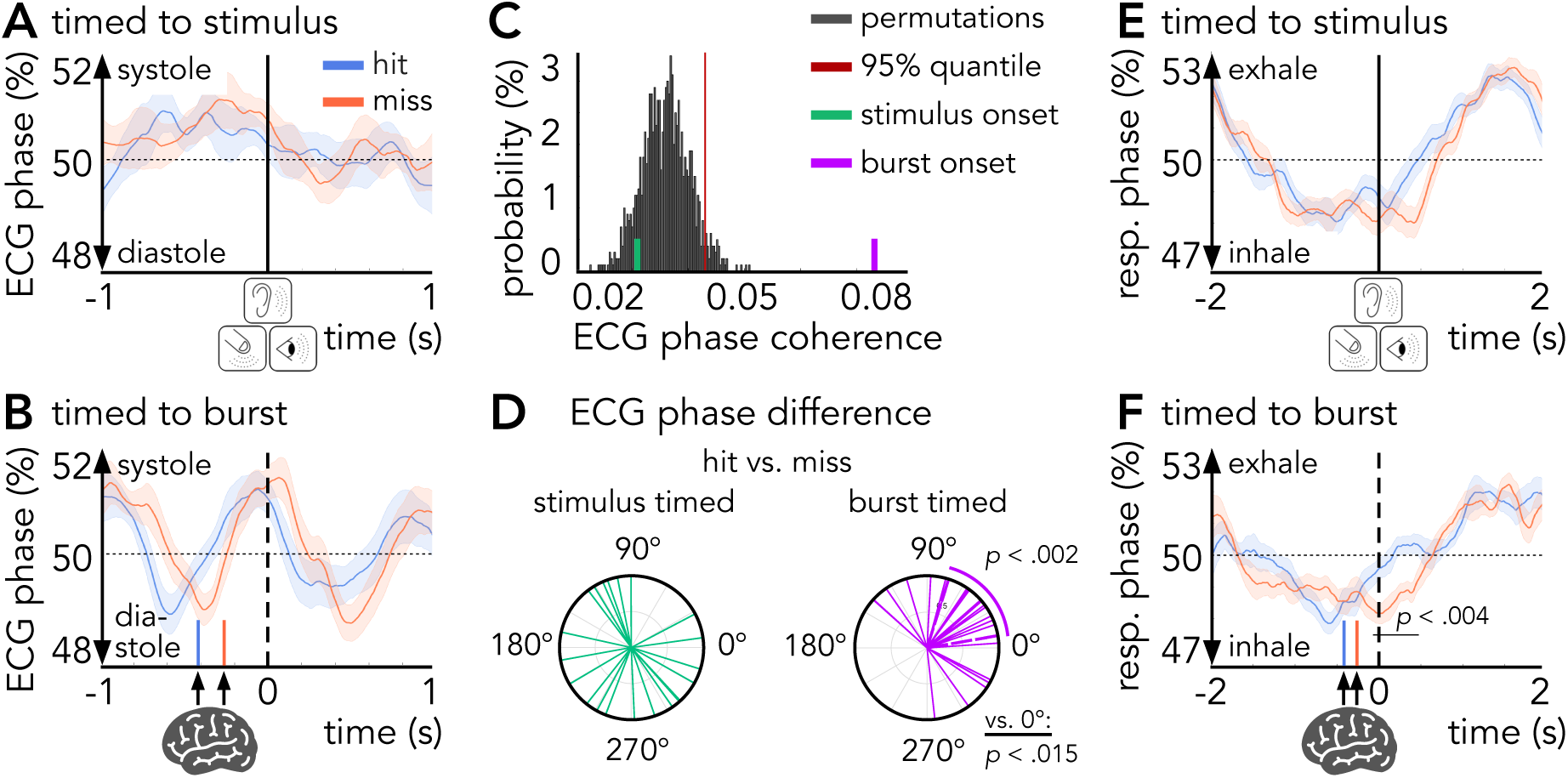
Cardiac and respiratory activity per perceptual outcome relative to stimuli and neural bursts **A** ECG phase locked to stimulus onset (straight vertical line) for all detected (blue) and missed trials (red). Increasing values show more systolic- and decreasing values more diastolic activity. Shaded error regions represent SEM for repeated-measures designs. **B** as in **A** but time-locked to burst onset and then shifted to the average onset times per participant and perceptual outcome (inset vertical lines) relative to stimulus onset (dotted vertical line). See Figure 3C for a schematic overview of the method. **C** Inter-trial ECG phase coherence at stimulus-(green) and burst onset (purple), and for 1000 random permutations. The red vertical line denotes the 95%-quantile of the permutation distribution. **D** ECG phase difference for hit- vs. miss-trials per participant for the stimulus-timed ECG (green) and the MEG-timed ECG (purple). Thick, straight lines show the mean phase value and dotted lines show the 95%-quantiles for the MEG-timed ECG. The stimulus-timed ECG data did not meet the criteria to calculate CIs. *P*-values denote significance levels for the inter-participant ECG phase coherence (inset circular segment) and for the phase difference from 0°. **E** and **F** as in **A** and **B**, respectively, but for respiration phase. Increasing values show more exhalation and decreasing values more inhalation. *P*-values denote significance levels for repeated-measures *t*-statistics (cluster-corrected).

## Discussion

### Conscious awareness is an “inside job” orchestrated by internal timing

The present work demonstrates that conscious perception is not determined by external stimulation alone, nor by ongoing brain activity in isolation, but by the active temporal alignment between internally generated neural states, bodily rhythms, and incoming sensory events. Across auditory, tactile, and visual modalities, perception depends on whether sensory input arrives at a moment when the brain had stabilized an internal reference frame - a process whose timing is intrinsically coupled to cardiac cycles and can be modulated through respiration. These findings reveal a supramodal organizing principle: conscious access across sensory systems is governed by internally structured timing processes that the observer partially regulates. Perception is thus orchestrated from within - shaped by self-generated brain-body dynamics that temporally gate when external inputs can enter awareness.

### Alpha bursts temporally organize perceptual readiness

By resolving alpha activity into temporally resolved bursts, we identify a mechanism that explains why identical stimuli may be detected or missed. Detected trials were characterized by earlier and more temporally clustered alpha bursts in pre-stimulus periods, indicating that an internal reference frame had stabilized before stimulus arrival — a prerequisite for perceptual stability in continuous experience.^52^ When bursts occurred later or were temporally dispersed, stimuli coincided with a less stable internal configuration and failed to gain conscious access. Perceptual outcome was thus determined not simply by the overall amount of alpha activity, but by when internally generated states reach stability relative to sensory input. These findings build on prior work^21,22^ linking higher pre-stimulus alpha power to misses, particularly in vision, which our power analysis replicated. Mean power differences have been interpreted in terms of fluctuations in overall cortical excitability and corresponding shifts in decision criterion within a signal detection framework.^53–55^ Our burst-based analysis refines this perspective by isolating the temporal structure of alpha activity. Whereas alpha power may reflect the overall level of ongoing activity, burst timing captures transient state transitions relative to this background. Because burst detection is defined with respect to local baseline activity, differences in tonic alpha power can influence burst estimates, such that power and burst measures need not vary in the same direction and may, under certain conditions, diverge. In this context, alpha power may index global excitability or response bias, whereas the timing of alpha bursts may reflect transient internal states that shape moment-to-moment perceptual sensitivity. The earlier and more temporally precise occurrence of bursts on detected trials suggests that these events mark internal transitions that facilitate access to sensory information, beyond baseline shifts in excitability alone. In this sense, bursts reflect the emergence of a perceptually ready state within ongoing brain activity. The supramodal consistency of these effects - shared timing, alpha-band specificity, and ipsilateral localization in premotor and prefrontal regions – argues against sensory-specific explanations and instead aligns with evidence for common supramodal neural signatures of conscious perception across sensory systems.^44^ Alpha bursts may index a higher-order, network-level control process through which the brain regulates excitation–inhibition balance to proactively configure itself for upcoming input. By stabilizing a reference frame in advance, internal dynamics become predictive: the observer shapes the temporal conditions under which incoming stimuli can be integrated into conscious experience.

### Alpha desynchronization reflects internal state transitions rather than stimulus onset

Traditionally, post-stimulus alpha desynchronization has been viewed as a marker of cortical engagement or stimulus-driven processing.^15,45,46^ Our findings motivate a reinterpretation: alpha desynchronization reflects the completion of an internally generated state transition rather than the initiation of stimulus processing. In this view, perception depends on how ongoing anticipatory dynamics—established before stimulus onset—evolve through and beyond stimulus arrival. This interpretation aligns naturally with classical findings of lateralized alpha power asymmetries in spatial attention paradigms, typically interpreted as inhibition of task-irrelevant space and indirect facilitation of contralateral processing.^19,20,56–58^ These spatially specific effects can be understood as an implementation of anticipatory state configurations, in which alpha dynamics prepare the system for expected input by structuring the allocation of neural resources across space. Consistent with this framework, burst-based analyses showed that predominantly ipsilateral pre-stimulus alpha activity indexes the formation of a temporally structured internal state, whereas post-stimulus desynchronization reflects its reconfiguration. On detected trials, this internally generated state resolves shortly after stimulus onset, allowing sensory input to be integrated into a newly configured perceptual regime. On missed trials, by contrast, alpha activity persists into the post-stimulus interval, indicating that the system remains in an internally oriented configuration despite external input. This state-transition perspective subsumes inhibitory accounts of alpha function^15,16,59,60^ within a broader temporal organization framework. Alpha dynamics regulate whether and when the system transitions from an internally oriented regime into a state receptive to incoming information. Importantly, this transition is not purely stimulus-driven but reflects the continuation of pre-stimulus expectancy dynamics into post-stimulus processing. Within this framework, desynchronization complements the burst findings: while bursts index the emergence and stabilization of a pre-stimulus reference state, desynchronization marks the completion of its transition as perception unfolds. Our burst findings replicate the classical lateralized alpha power pattern across auditory, tactile, and visual modalities, but show that it reflects a more general mechanism—complementing, rather than replacing, spatially selective interpretations—by indexing the reconfiguration of a supramodal predcitive internal state. Conceptually, alpha desynchronization thus marks a shift in the system’s semantic orientation—from a state of anticipation to one of perceptual engagement—consistent with accounts linking alpha dynamics to controlled access to internal representations and large-scale network organization.^61–63^ In this sense, desynchronization is not the onset of perception but the signature of a completed internal transition that renders perception possible.

### Brain–body coupling as a temporal scaffold

Because alpha bursts provide temporally resolved markers of internally stabilized neural states, they enable the identification of precise temporal alignment between brain activity and peripheral physiological rhythms. Burst timing was systematically coordinated with cardiac and respiratory cycles, suggesting that ongoing bodily signals contribute to the temporal organization of perceptual readiness. In contrast to the typically found desynchronization after external stimuli, neural alpha burst activity increased when aligned to internally generated cardiac and respiratory events. Reversing the analytical frame revealed consistent phase relationships between cardiac and respiratory signals and burst timing—dependencies that are not observable in conventional stimulus-centered analyses. Together, these findings indicate a structured temporal coupling between neural and physiological systems, in which brain dynamics and bodily rhythms form coordinated timing relationships. This pattern is consistent with proposals that brain and bodily oscillations form a hierarchical frequency architecture, in which cross-system timing relationships provide a structural scaffold for neural processing and perception^64,65^, and may reflect a general principle of large-scale neural organization by cross-frequency coordination.^66^ Importantly, the timing of this coupling relative to stimulus onset predicted perceptual outcomes. These results are compatible with theoretical accounts proposing that perception is embedded within an internally generated reference frame shaped by brain–body interactions.^67^ Integrating recent findings further suggests that alpha oscillatory timing not only gates sensory input but also structures internal representations of the body, linking self-related processing to perceptual access.^68^ In this view, the sense of an “observer” arises from the alignment of neural and physiological dynamics prior to and during stimulus processing, rather than from stimulus-driven processes alone. Together, these results suggest that brain–body timing provides an endogenous temporal structure that constrains when sensory inputs become accessible to awareness.

Importantly, our findings do not imply a linear causal chain from heart or respiration to perception but support a model in which neural and physiological rhythms form a coupled timing system. In this view, bodily rhythms do not drive perception directly; they constrain the temporal windows in which internal neural states can form and dissolve, creating an internally generated temporal scaffold for awareness. Several physiological mechanisms may underpin the observed brain–body interactions. The heart-brain connections are likely an intrinsic feature, given their presence both during the task and at rest. Prior studies^33–35,49,50^ indicate that afferent signals travel via the vagus and glossopharyngeal nerves and enter the brain through brainstem and thalamic nuclei, ultimately reaching cortical targets in cingulate, motor, and frontal regions — a pattern confirmed by our source reconstructions. However, we cannot rule out other global neuromodulatory influences (e.g., ephaptic coupling^69,70^ or intracranial pressure fluctuations^71,72^).^73^ Whereas cardiac cycles contribute a predictable temporal structure, respiratory cycles may provide an adaptable temporal mechanism for modulating neuro–cardio timing and cortical excitability. Participants’ breathing behavior was systematically related to task demands, suggesting that the timing of internal physiological states can adapt to ongoing cognitive context. When the brain is stabilized by alpha control networks, when the heart enters its activity-silent state after systole, and when exhalation-related changes in autonomic activity delay subsequent neuro–cardio interactions, stimuli may arrive at moments of reduced internal noise. This coordinated neuro–cardio–respiratory timing may thus define an internally configured perceptual regime that shapes optimal conditions for conscious access. Real-time triggering of stimuli at defined physiological phases and/or aligned with neural bursts could provide an ecologically valid test of these dynamics. Computational modeling may further formalize how recurrent brain–body feedback loops generate temporally structured perceptual states. Rather than assuming linear causality, these models should emphasize cyclic, self-organizing dynamics (e.g., attractor networks^74,75^ or manifolds^76,77^), in which neural and physiological rhythms jointly define internally stabilized perceptual states.

### Limitations and future directions

Several limitations should be considered when interpreting the present findings. First, although we employed multiple complementary preprocessing and validation approaches to mitigate cardiac field artifacts (see Methods and Supplementary Fig. 2), their influence cannot be fully excluded given the close temporal coupling between cardiac and neural signals. Importantly, increasing levels of artifact mitigation reduced correlations between corrected MEG and ECG timeseries while preserving similarity with the original MEG signal in both empirical data and simulations. This pattern indicates progressive attenuation of cardiac field artifacts with minimal distortion of the underlying neural signal. Moreover, control analyses showed that burst timing was not zero-phase-locked to the QRS complex, arguing against a primary contribution of residual cardiac field artifacts. Second, the identification of oscillatory bursts depends on the specific definition of transient activity, and ongoing debate remains regarding whether neural oscillations are best characterized as sustained or burst-like. To address this, we implemented multiple burst detection approaches, including methods based on local amplitude thresholds as well as a waveform-based approach that does not primarily rely on amplitude normalization (see Methods, Fig. 3A and Supplementary Fig. 1). The convergence of results across these methods indicates that the observed effects are not driven by a particular detection algorithm or by high-amplitude events alone. Third, all stimuli were presented in the left visual field, which limits the ability to fully disentangle ipsilateral effects from more general spatial asymmetries. While this design ensured consistent spatial attention demands, future work should test whether the observed effects generalize across hemifields.

### Summary

Our findings demonstrate that conscious perception across sensory systems is shaped by temporally precise, internally generated brain states coordinated with physiological rhythms. Alpha bursts mark transient stabilization events that establish perceptual readiness, whereas alpha desynchronization signals transitions between internal states. Cardiac and respiratory cycles temporally structure these neural dynamics, revealing a multiscale temporal architecture that organizes perceptual processing in time. Conscious awareness may therefore arise from the coordinated progression of neural and physiological states that gates access of sensory inputs to consciousness.

## Methods

### Ethics statement

All procedures were approved by the ethics committee of the University of Salzburg.

### Participants

Twenty-one participants (14 female; mean age ± *SD*, 26.3 ± 5.2 years) took part in the experiment. Respiration data was recorded from a subset of fourteen participants. All participants were right-handed, had normal or corrected-to-normal vision, normal hearing, reported no history of neurological or psychiatric disorders, gave written informed consent before the experiment, and received a monetary compensation or course credits.

### Apparatus

#### MEG, ECG and respiration data acquisition

Electrophysiological brain activity was recorded with a whole-head MEG system with 102 magnetometers and 204 planar gradiometers (Neuromag306; Elekta), sampled at 1000 Hz in a magnetically shielded room. For each participant, a head frame coordinate reference was defined before the experiment by digitizing the cardinal points of the head (nasion and left and right preauricular points), the location of five head position indicator coils, and a minimum of 200 other head shape samples (3Space Fastrack; Polhemus). The head position within the MEG helmet (relative to the head position indicator coils and the MEG sensors) was controlled before each block to ensure that no large movements occurred during the data acquisition. ECG was recorded using two standard electrodes, one placed just beneath the left clavicle at the Infraclavicular fossa and the other directly under the midpoint of the left armpit at the midaxillary line. The abdominal respiratory movements were measured with a breathing belt (Respiration Belt MR; Brain Products).

#### Stimulus presentation

Stimuli were generated using MATLAB 8.6 (The MathWorks) and Psychophysics Toolbox (PTB), version 3^78,79^ using the objective PTB extension.^80^ A DLP projector (PROPixx; VPixx Technologies) showed the visual stimuli at a refresh rate of 120 Hz onto a translucent screen (26 horizontal x 14 vertical degrees of visual angle [DVA]) in front of the participant (viewing distance, 120 cm) within the dimly lit, magnetically shielded MEG room. Visual stimuli were shown to the left side relative to the fixation cross, pure-tone sounds were presented through an earplug placed in the participants’ left ear and tactile stimulation was delivered through independently vibrating tips on a vibrotactile stimulator attached to four fingers of the participants’ left hand (Brain Gauge Research Edition; Cortical Metrics). Stimulus timing was controlled with a data and video processing peripheral (DATAPixx; Vpixx Technologies).

### Stimuli and experimental design

Before the experiment, all participants completed 20 practice trials at high stimulus intensities for each modality to familiarize with the experimental setup and the response procedures. Each trial began with the central presentation of a black fixation cross with a size of 1.5 DVA on a gray background, which remained on the screen until the response period. After a temporally jittered pre-stimulus period for 1-2 s duration, an auditory, tactile or visual stimulus was presented for 50 ms (see Fig. 1A). For the present analyses, stimuli were pooled within modality. The auditory stimuli consisted of pure sine-wave tones at 200, 431, 928 or 2000 Hz delivered to the left ear. The visual stimuli were Gabor-patches tilted to either vertical, horizontal or +/- 45° rotated orientations (size: 4 DVA, spatial frequency: 0.7x10^-3^ cycles per DVA, *SD* of the Gaussian envelope: 0.8 DVA) shifted horizontally to the left from center for 4 DVA. For tactile stimulation, we delivered vibrotactile stimuli at 30 Hz to either the left index-, middle-, ring- or little finger. On each near-threshold trial, the respective stimulus intensity was selected by an adaptive algorithm based on Bayesian inference to reach approximately equal proportions of detected vs. missed reports.^81^ The mean stimulus intensities ± *SD* per modality were for auditory stimuli -64 ± 3.7 dB, for tactile stimuli 13.7 ± 2.6 % of maximum stimulator output, and for visual stimuli 5.5 ± 2.2 % contrast. Moreover, we used clearly perceptible sounds of -10 dB, vibrations with 100% stimulator output and Gabors with 100% contrast for supra-threshold, high-signal trials, whereas no stimuli were presented on no-signal, catch trials. Stimulus presentation was followed by a jittered post-stimulus period for 0.5-1 s duration and a response period for a maximum duration of 5 s, in which a central question mark (font size: 20) alongside a blue circle to the left and a yellow circle to the right of 1.3 DVA size, indicating the respective response buttons, were shown. Below the circles, we presented either the word “yes” or “no”. The response mapping between the blue and yellow buttons and the yes-no responses was randomized over trials. Each button press was followed by a wait period for 500 ms duration before the next trial began. The three modalities were run in counterbalanced blocks across participants. We used four blocks each comprising 120 trials per modality (i.e. twelve blocks total) with the different stimuli per modality balanced and randomly intermixed within a block. Ten percent of trials were dedicated to high signal- and catch trials, respectively, per block, and the remaining 80% trials were near-threshold trials. For one participant, data from only 11 blocks was available due to technical difficulties. Each block was about 7-8 minutes in duration and the entire experiment lasted approximately 1.5-2 hours with small breaks between blocks.

### Behavioral data analysis

Behavioral performance was quantified in terms of percent yes-responses for each modality, each stimulus per modality, and per near-threshold, high signal and no signal trial types (see Fig. 1B). Further, we performed a signal-detection analysis to estimate sensitivity and response bias by calculating d-prime (d’) and criterium (c) for each modality. All error bars (for behavioral, MEG, ECG and respiration measures) show the SEM for repeated measures when appropriate. The mean between conditions was subtracted from the data in each condition before calculating the SE. The resulting error estimate was bias corrected by the number of conditions (M), this means it was multiplied by ✓(M/(M-1)).^82^

### MEG data analysis

#### Data preprocessing

The data were analyzed using custom-built MATLAB code (MATLAB 9.13; The MathWorks) and the FieldTrip toolbox.^83^ We used a signal space separation algorithm implemented in the Maxfilter program (version 2.2.15) provided by the MEG manufacturer to attenuate external noise from the MEG signal (including signals arising from volume conduction in the participants’ body, but also 16.6 Hz, which is the alternating current used by the railway electrification systems in Austria, and 50 Hz plus harmonics, which is line noise in Europe) and to compensate for small head movements during the experiment by realigning the data to a common standard head position based on the measured head position at the beginning of each block. Continuous data were high-pass filtered at 0.5 Hz with a FIRWS filter (2 Hz transition width), low-pass filtered at 99 Hz, and band-stop filtered between 49-51 Hz with a two-pass Butterworth filter (order 4), applied in the forward and reverse directions. Then, MEG data were segmented from -2 s to +2 s relative to stimulus onset. Unless otherwise indicated, all data show MEG activity on combined gradiometer channels. Data from the horizontal and vertical component of the planar gradient were combined via their vector sum. The effects on magnetometer channels were qualitatively and quantitatively similar.

#### Artifact rejection

A set of summary statistics (variance, maximum absolute amplitude, maximum z value) was used to detect statistical outliers of trials and channels in the datasets. These trials and channels were removed from each dataset (semiautomatic artifact rejection). Moreover, data containing squid jump artifacts were automatically detected and removed. On average, 5 channels (2.3 *SD*) and 4.6 % of the trials (4.9 % *SD*) were rejected. The rejected channels were interpolated with the nearest-neighbors approach for sensor-level analysis. Interpolated channels were not used for source reconstruction and independent component analysis.

#### Independent component analysis

Independent component analysis (ICA) is the best-practice method to reduce cardiac field artifacts resulting from volume conduction (see for example ^36,38,41,84^). Before performing ICA, data were down-sampled to 250 Hz to speed up computations. ICA was run using the infomax algorithm (method ‘runica’) implemented in EEGLAB.^85^ The component time courses for each trial were then correlated with the corresponding ECG, as well as vertical and horizontal electro-oculography (EOG) time courses, and the mean absolute correlation coefficients per component were computed. Subsequently, components exceeding a threshold of two times the median correlation divided by 0.6745 were removed (note: the threshold definition is based on the median absolute deviation^86^, see the section “detection of oscillatory bursts” below for details). On average, this led to the removal of 6.2 components (2.8 *SD*).

#### Removal of cardiac field artifacts for real and simulated data

On top of ICA, we used an additional method to correct for cardiac field artifacts (CFA). We time-locked each data segment to the time point of the R-peak ±2 s, computed the across-trial average field and subtracted it from each individual data segment. This method was highly effective in removing CFA in our data (see Supplementary Fig. 2A) and in simulation work (see Supplementary Fig. 2B). For the simulations, we artificially introduced a dipole corresponding to a typical CFA into original, ICA-cleaned MEG data (see above) and estimated the impact of different artifact corrections methods (i.e. no cleaning, ICA, CFA removal) by correlating the corrected data with the simulated ECG, and with the original MEG before the CFA-dipole was added. The artifical ECG signal was simulated using Python (Python version 3.13^87^) and the function ‘ecg_simulate’ implemented in the NeuroKit2 toolbox (NeuroKit version 0.2.0^88^; noise settings = 0.01, heart_rate = 65, heart_rate_std = 4, method = ’ecgsyn’). It was then added to the previously cleaned MEG recording using a modified version of the MNE Python function ‘add_ecg’ (MNE Python version v1.9.0^89^). The ECG dipole location and orientation matched with typical CFA in MEG data.^84^ Further, expected measurement noise (pink noise) was added to the the simulated ECG signal (prior to dipole simulation) and to the MEG signal (after dipole simulation). The CFA removal method resulted in the lowest correlations between the corrected MEG data and the simulated ECG timeseries, while maintaining the highest similarity with the original MEG data. This pattern suggests that the method effectively attenuates cardiac field artifacts with minimal distortion of the underlying neural signal.

#### Stimulus-evoked fields

Before computing the evoked fields, MEG data were low-pass filtered at 30 Hz. Evoked fields were then computed as the across-trial average MEG activity per modality and trial type and baseline corrected for the interval from -0.2 s to 0 s. The time courses shown in Fig. 1C display the average evoked fields over contralateral auditory, visual and somatosensory areas (see inset topographies in Fig. 1C). The gradiometer topographies shown in Fig. 1C show the average evoked fields from 0 s to +0.5 s for detected trials per modality vs. baseline.

#### Time–frequency representations

Before calculating the time-frequency representations (TFR), we subtracted the trial average from each single trial, to obtain induced activity without the contribution from stimulus-evoked components. TFRs were calculated on single-trial data using Morlet-Wavelets applied to short sliding time windows in steps of 10 ms in the time interval between -1.5 s and +1.5 s relative to stimulus onset and in the frequency range between 5 Hz and 50 Hz. We used a frequency-dependent window width of 5 cycles per frequency. The squared absolute value gave the signal power for each MEG sensor across different frequencies and time points. The time-frequency maps shown in Fig. 2 display the relative power change in % from the baseline period from -1.5 s to -1 s ((MEG power – BL) / ((MEG power + BL)/2)) on a representative MEG gradiometer sensor (black dot on sensor-level topographies in Fig. 2) separately for the baseline, detected and missed, near-threshold trials per modality.

#### Detection of oscillatory bursts

We implemented a thresholding algorithm for oscillation burst detection^27,31^ (for review see^30^), derived from automatic spike-detection methods (adapted from^90^). First, the MEG data were band-pass filtered between 8-13 Hz to exclusively extract alpha frequency band activity. Then, we applied the Hilbert-transform on the band-limited data and computed its amplitude envelope. Subsequently, burst activity was detected when the MEG amplitude exceeded the median amplitude of the respective trial between -2 s to +2 s divided by 0.6745 for at least three alpha cycles (see Equation 1 and Fig. 3A). We selected this method for the main analyses because it equalized overall burst counts between detected and missed trials (see Supplementary Fig. 1), allowing temporal differences to be interpreted as genuine timing effects rather than consequences of burst prevalence.

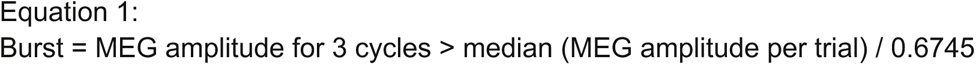

To define a robust amplitude threshold, we made use of the statistical concept of the probable error, which defines the half-range interval around a distribution’s central point. For symmetrical distributions, it is equivalent to the interquartile range or the median absolute deviation.^86^ The probable error can be expressed in terms of SDs when scaled with a constant factor that depends on the assumed distribution (i.e. this factor refers to the 75% quantile for a normal distribution with <l^-1^(0.75) = 0.6745). It is a more robust measure of variability compared with the SD because it is more resilient to outliers (i.e., it remains largely constant across different burst rate regimens and amplitudes). Moreover, because the variability per trial and channel is estimated across the MEG signal of the entire trial, it is independent from trial-specific baseline periods. The bursting analysis results in sparse time courses for each trial and channel, which separate the occurrence of a suprathreshold burst event from non-burst periods. Averaging the sparse burst activity time courses over detected and missed, near-threshold trials gives the burst rate in % shown in the time courses in Fig. 4 and 5 for a representative MEG gradiometer channel (black dot on topographies). Additionally, single-trial metrics of burst activity timing were computed by extracting the burst onset- and offset times within the interval -1 s to +1 s relative to stimulus onset. The average burst onset times are shown as inset circles in Fig. 4 and 5. The leftmost topographies shown in Fig. 4 and 5 display the proportion of trials containing a burst in the interval between -1s to +1s. The other topographies show dependent-samples *t*-statistics between detected vs. missed, near-threshold trials for the burst onset times and the burst rates (see Statistical analysis). Further, we computed the burst durations and their timing variability estimated by the SD of the onset times across trials (shown in Supplementary Fig. 1). The detection of transient bursts of oscillatory brain activity, instead of the more traditional power analysis, was motivated by recent reports from neurophysiology showing that oscillatory burst events underlie many cognitive and motor operations (e.g. ^27,91–93^). These observations have sparked a debate on principles about the sustained versus transient nature of oscillatory dynamics in the brain (for discussion see^29^). The key aspect is that, in many cases, sustained oscillatory activity represents an analysis artifact from averaging over trials, and it may instead be better captured by transient, high-signal burst events that happen at different rates and times, and with different durations from trial to trial. Moreover, the burst-analysis strategy makes much sense from a computational modelling perspective, because burst events are thought to represent supra-threshold reverberations of latent states embedded in network connections, in line with predictions from attractor network theories.^74,75,94,95^

#### Sensitivity analysis of oscillatory burst detection

Because burst detection relies on thresholding relative to background activity, differences in overall alpha power between conditions (e.g., hits vs. misses) could in principle bias burst estimates. To address this, we systematically evaluated five complementary burst detection approaches that varied in their dependence on baseline power (Fig. 3A). Across all methods, results were qualitatively and quantitatively consistent (Supplementary Fig. 1). The primary method (Equation 1), used in the main analysis, estimated baseline power within each trial, providing a sensitive measure of the relative prominence of transient events but potentially reflecting condition-specific power differences. To reduce this dependency, we implemented three alternative threshold-based approaches that progressively relaxed the dependence on trial-specific amplitude estimates. First, thresholds were derived from the pre-stimulus interval (−1.5 to −1 s; Equation 2), reducing within-trial variability. Second, a local estimate based on a ±5-trial sliding window (Equation 3) further reduced sensitivity to trial-by-trial fluctuations. Third, a condition-level threshold was computed by pooling detected and missed trials (Equation 4), thereby eliminating condition-specific baseline shifts. Together, these approaches ensured that burst detection was not driven by systematic amplitude differences between conditions. In all cases, bursts were defined as periods in which the signal exceeded the respective threshold for at least three alpha cycles.

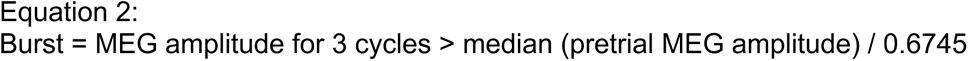

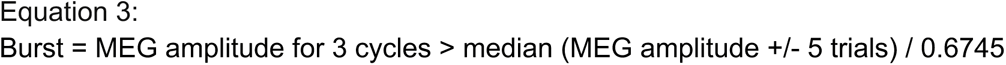

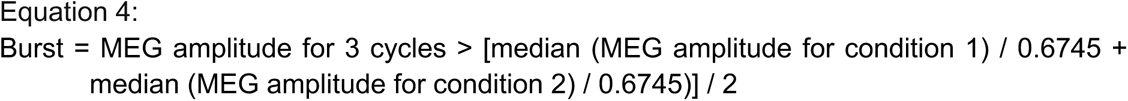

Finally, we implemented a waveform-based burst detection method (included in the cycle-by-cycle toolbox^96^) that does not rely on amplitude thresholds. Instead, it identifies bursts based on the temporal structure of the signal on a cycle-by-cycle basis, providing a threshold-free estimate of oscillatory burst events and capturing non-sinusoidal features of the MEG signal (Fig. 3A, Supplementary Fig. 1). First, data was bandpass filtered in a broader range between 4 and 17 Hz. Then, burst activity was detected when the MEG signal waveform satisfied a set of parameters with respect to the amplitude consistency of adjacent cycles (> 0.4), the average amplitude relative to all other cycles (amplitude fraction, > 0.2), the period consistency of adjacent cycles (> 0.4) as well as the monotonicity of the rise and decay flanks of the cycles (>0.8). The parameter settings were selected based on previous work.^32^ The frequencies of the burst cycles were estimated based on the time between consecutive troughs and the frequency range of interest was set to 8 – 13 Hz. As for the other methods, burst activity was defined as periods during which the MEG signal met these parameters for at least three consecutive alpha cycles (Equation 5). Waveform analysis was performed for each planar gradiometer separately to preserve the required polarity information. We used the data from the vertical gradiometer channel for single-trial analysis. The data from the horizontal gradiometer was similar on average, but noiser on the level of individual trials. Data from both gradiometers were averaged across trials and then combined to plot topographies.

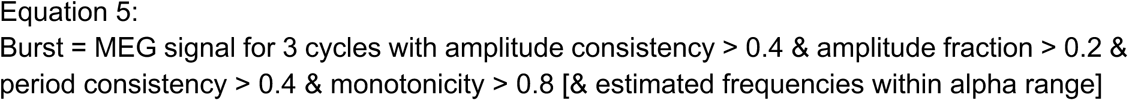

The convergence of the findings across all methods indicates that burst effects are not driven by the specific choice of baseline definition.

#### Source reconstruction

To localize the cortical generators of the observed sensor-level effects, we used source reconstruction based on a spatial-filtering algorithm (i.e. beamforming). A structural MRI was available for 8 of 21 participants, for which we co-registered the brain surface from their individual segmented MRIs with a single-shell head model.^97^ For the remaining participants, we obtained the canonical cortical anatomy from the affine transformation of an MNI template brain (brainweb.bic.mni.mcgill.ca/brainweb/) to the participants’ digitized head shape. This hybrid approach is standard in group-level MEG analyses when individual MRIs are unavailable. Then, we created MNI-aligned grids in each participant’s individual headspace. To this end, each participant’s head shape was warped to the MNI template brain, and the inverse of the transformation matrix was applied onto an MNI template grid with 889 grid points and, on average, 1.5 cm spacing. With this method, we achieved a consistent mapping of the spatial positions of grid points across participants and to the MNI template. Single-trial, MEG-sensor time courses were projected into source space (individual MNI-aligned grids) using a linear constrained minimum variance beamformer algorithm (lcmv).^98^ We defined a common spatial filter based on the covariance matrix of the band-pass filtered signal (cutoff frequencies, 1–30 Hz). The covariance window depended on the locus of the effect as expected from sensor-level analysis (-0.5s to +0.5s relative to stimulus onset). The beamformer filters were computed using the broadband single-trial data. Both the information from the magnetometer and planar gradiometer sensors systems were used. Oscillatory burst detection (see above) was performed on the single-trial, source-space data and the corresponding alpha burst rates were calculated. Next, we contrasted the average burst onset times and the burst rates averaged separately over the pre-stimulus (-1s to 0s) and the post-stimulus intervals (0s to +1s) between detected vs. missed, near-threshold trials using dependent-samples *t*-statistics and correcting for multiple comparisons at multiple voxels with a cluster-based permutation approach (see Statistical analysis). The resulting difference maps and the proportion of trials containing a burst in the interval between -1s to +1s (masked at the upper 10% voxels) were interpolated onto a standard MNI brain (Fig. 5G). Anatomical structures corresponding to localized sources in MNI coordinates were found using the software MRIcron (https://www.nitrc.org/projects/mricron/).^99^

#### R-peak timed and respiration-peak timed analysis

To test whether burst timing aligned with internal physiology, we re-expressed single-trial burst activity time series relative to the last R-peak or respiration peak preceding stimulus onset (−1 s to 0 s) for each trial and participant. On average, R-peaks were detected in 98.2% of trials (SD = 3.9%) and respiration peaks in 35% of trials (SD = 8.4%) within this interval (see section *ECG and respiration data analysis* for peak detection procedures). Burst rates and single-trial burst timing metrics were then computed in this cardiac- or respiration-centered reference frame, separately for detected and missed near-threshold trials for each participant. Finally, the resulting time series were projected back into stimulus time by shifting them according to each participant’s mean pre-stimulus R-peak or respiration peak times, thereby restoring a common stimulus-centered time axis across trials. This analysis preserves the original temporal relationship to stimulus onset while revealing burst dynamics relative to intrinsic cardiac and respiratory cycles, thereby enabling direct comparison with stimulus-locked analyses (see Fig. 5A–C). Conceptually, this procedure corresponds to briefly viewing neural activity from the perspective of cardiac or respiratory events before returning it to a common stimulus-centered timeline (see Fig. 3B).

#### Resting state data

Each participant’s resting state activity was recorded after the experiment with their eyes opened and fixated at a central fixation cross for 5 minutes. Unless otherwise indicated, all preprocessing and analysis steps for the resting state recording were identical to the task data described above (including the methods to correct for CFA, the calculation of TFRs and oscillatory bursts, and the source analysis). As shown in Supplementary Fig. 3, we detected all R-peaks during the resting state recording for each participant, time-locked and segmented the MEG data to -1.5s to +1.5s around each R-peak and calculated the TFRs and oscillation burst rates. For comparison, we proceeded in the same way for the task data by referencing it to the last R-peak in the interval from -1s to 0s relative to stimulus onset on each trial. Supplementary Fig. 3A and C shows the R-peak locked power increase in % signal change from a neutral baseline period, in which no visible heartbeat-evoked activity was present (-1s to -0.5s for the task data, -1.5s to -1s for the resting state data) averaged over central gradiometer channels. Supplementary Fig. 3B and D shows the burst rate change after vs. before R-peaks in % signal change for the task data and the resting state data, respectively, interpolated onto a standard MNI brain (masked at the upper 10% voxels). For this analysis, we explicitly excluded the time periods around the R-peaks (-0.1s to +0.1s) and computed the pre-R-peak period from -0.3s to -0.1s and the post-R-peak period from +0.1s to +0.3s, to reduce any volume conduction effects from cardiac field artifacts.

#### Summary of methods to mitigate cardiac field artifacts

Because cardiac field artifacts are an important confound, we summarize all employed methods to mitigate them in our data. First, the signal space separation algorithm implemented in the Maxfilter program removes environmental noise including from the participants’ body from the MEG signal. Second, we focused the main analysis on the gradiometer MEG channels, which are known to be more sensitive to activity generators close to the cortical surface and are less influenced by deeper sources. Third, the lcmv-beamforming algorithm used for source reconstruction adds a further spatial filtering step concentrating the analysis more towards cortical rather than bodily origins. Fourth for some analyses, we explicitly excluded the time periods around the R-peak, while for other analyses we show the full temporal evolution showing that the MEG effects, while located in the temporal vicinity of the R-peak, do not temporally coincide with it, as it would be expected from volume conduction effects. Fifth, we used ICA to reduce CFAs (in accordance with prior work^36,38,41,84^). Sixth, we implemented a direct removal approach of CFAs by subtracting the average R-peak locked activity per participant from each trial. This method was effective in removing CFAs in our data and in simulation work (see Supplementary Fig. 2).

### ECG and respiration data analysis

The initial data preprocessing of the ECG and respiratory signals was implemented using the Python-based NeuroKit2 toolbox (Python version 3.13^87^; NeuroKit version 0.2.0^88^). It features data cleaning and the automatic extraction of the ECG- and respiratory signal components, namely the QRS-complex and the T- and P-waves of the typical ECG, and the respiration peaks and troughs. Subsequent analyses were run using custom-built MATLAB code (MATLAB 9.13; The MathWorks) and the CircStats toolbox (Version 1.21.0.0^100^). The ECG data from one participant was too noisy and therefore excluded from further analysis of the ECG time course. However, we recovered the R-peak timestamps for this participant by thresholding their raw ECG time series with 5*above its median activity. This worked well, due to the high signal-to-noise ratio of the R-peak. For all participants, the physiological signal time courses were segmented from -2 s to +2 s relative to stimulus onset and smoothed with a Savitzky-Golay-filter with a kernel-width of 150 ms to reduce temporal noise. For the main analysis, we focused on the ventricular ECG phase. The data for the atrial ECG was quantitatively and qualitatively similar. The ventricular ECG systole is defined from the R-peak to the end of the T-wave entering the diastolic phase afterwards until the next R-peak. Figure 6A and B shows the ventricular ECG phase in %, averaged separately over detected and missed, near-threshold trials, such that values above 50% indicate a greater probability of systolic activity and below 50% more diastolic activity. Figure 6A shows the data segments time-locked to stimulus onset. By contrast, Figure 6B shows the same data but time-locked and averaged to burst onsets at a representative gradiometer channel and then projected back to the mean burst onset times per participant and perceptual outcome, to keep the appropriate temporal reference with stimulus onset (see Fig. 3C). Burst onsets in the interval from -2 s to +2 s were included, to account for the slower temporal dynamics of the physiological data. The ECG phase at burst- and stimulus onset was computed by extracting the R-peak times before and after each onset per trial and then translating the respective onset times into a phase value within the cardiac cycle using the formula in equation 6 (see^38^ for a similar approach).

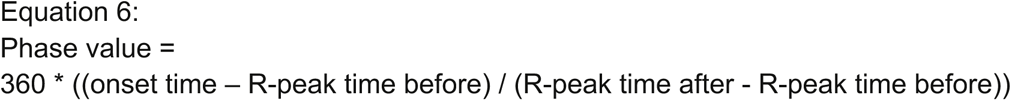

The ECG phase coherence at burst- and stimulus onset shown in Figure 6C was calculated by computing the mean resultant vector length across trials. It was compared against a random permutation reference distribution of ECG phase coherence values for statistical inference (see section Statistical analysis below). The polar histograms shown in Figure 6D display the ECG phase difference between detected vs. missed, near-threshold trials (i.e. their circular distance) per participant separately for the stimulus-timed ECG and the neural burst-timed ECG. The burst-timed ECG phases were computed by translating the burst onset times relative to stimulus onset into a phase value based on each participant’s individual cardiac cycle period and then shifting the original ECG phase values at burst onset by this amount (similar to Figure 6B). We equalized any discrepancies in trial numbers between detected and missed trials by drawing a random subset from the condition with more trials, which was equal to the trial number of the condition with lesser trials. This process was repeated 1000 times and the average phase value across the subsets gave the mean phase value for that condition. Supplementary Fig. 4A and C displays the ECG-phase time-locked to all burst onsets (without shifting them back relative to stimulus onset) separately for the task data and the resting state data. Supplementary Fig. 4B and D shows ECG phase coherence values at burst onset and its corresponding random permutation reference distributions separately for the task data (same as Fig. 6C) and the resting state data. Figure 6E and F show the respiratory phase completion in % separately for detected and missed, near-threshold trials. It displays the temporal evolution of the four-phasic respiratory signal, such that signal increases indicate exhalation periods and decreases signify inhalation periods. Constant, horizontal signals denote periods, in which the participants hold their breath at maximum inhalation below 50% and at maximum exhalation above 50%. Figure 6E shows the respiration signal time-locked to stimulus onset, whereas Figure 6F shows the same data but time-locked and averaged to the neural burst onsets and shifted back to the average burst onset times per participant and perceptual outcome. Supplementary Fig. 5 displays the raw ECG- and respiratory signals time-locked to all respiration peaks or R-peaks, respectively, separately for the task data and the resting state data. The ECG phase coherence values at the respiration peaks and its corresponding random permutation reference distributions, shown in Supplementary Fig. 5C and F, were calculated in the same way as described above for the burst onset times.

### Statistical analysis

#### Cluster-based permutation statistics

We used a nonparametric, cluster-based permutation approach to correct for multiple statistical tests at multiple channels and/or time samples where appropriate.^101^ The nonparametric, cluster-based permutation procedure controls for the Type I error accumulation arising from multiple statistical comparisons. First, channel-temporal clusters of adjacent suprathreshold effects (dependent samples *t*-statistics exceeding *p*< .05, two-sided) were identified. The *t*-values within a connected cluster were summed up as a cluster-level statistic. Then, random permutations of the data were drawn by exchanging the data between conditions within the participants. After each permutation run, the maximum cluster-level statistic was recorded, generating a reference distribution of cluster-level statistics (approximated with a Monte Carlo procedure of 1000 permutations). The proportion of values in the corresponding reference distribution that exceeded the observed cluster statistic yielded an estimated cluster-level *p*-value (two-sided), which is corrected for multiple comparisons. It was used to correct for multiple tests at multiple channels for pre-stimulus (averaged between -1s to 0s) and post-stimulus (averaged between 0s to 1s) alpha-band power (averaged between 8-13 Hz) between detected vs. missed, near-threshold trials shown in Fig. 2. Then, it was used to correct for multiple channels, at which the difference in burst onset times between conditions was tested (shown in Fig. 4 and 5). Moreover, it was used to correct for multiple channel-time samples for the pre-stimulus (-1s to 0s) and post-stimulus (0s to 1s) burst rate differences (Fig. 4 and 5). It was further used for source-level statistics to correct for multiple voxels, at which the difference in burst onset times and the pre-stimulus (averaged between -1s to 0s) and post-stimulus (averaged between 0s to 1s) burst rates was tested (Fig. 5G). Finally, we employed it for the respiratory signals shown in Figure 6E and F to correct for multiple time samples within the interval between -1s to +1s relative to stimulus onset.

#### Bonferroni-correction for dependent-samples t-statistics

We opted for the Bonferroni-method to correct for multiple comparisons at multiple channels for the dependent-samples *t*-tests between pre-stimulus (averaged between -1s to 0s) and post-stimulus (averaged between 0s to 1s) alpha-band power (averaged between 8-13 Hz) vs. baseline (averaged between -1.5s to -1s), in order to set a common threshold for masking the sensor topographies shown in Fig. 2 and thus establish better comparability across the different modalities. We used the Bonferroni-method for the same reason to display the topographies for the different methods used in the sensitivity analysis (Supplementary Fig. 1). We decided for one-sided test statistics for the activation vs. baseline comparisons because of the clear directional hypothesis. We used two-sided tests between detected vs. missed trials for the sensitivity analysis.

#### Circular statistics for ECG phase

We used the CircStats toolbox for circular statistics (Version 1.21.0.0^100^). The consistency of the ECG phase across participants at burst- and stimulus onset, and for the circular distance between detected vs. missed trials was examined with a Rayleigh test. The difference in the ECG phase values between detected vs. missed trials, as well as between burst onsets times and R-peak times was investigated with an m-test testing whether the circular distance between the average phase values per condition across participants was different from 0°.

#### Permutation test for ECG phase

The consistency of ECG phase at burst onsets and respiration peaks across trials was assessed against a surrogate distribution obtained by breaking the trial-wise correspondence between neural and physiological signals. Specifically, burst onset times (or respiration peak times) were randomly permuted across trials while preserving the ECG signal within each trial, thereby destroying trial-wise alignment between neural events and cardiac phase. Since there is no trial structure for the resting state recordings, we calculated the reference distributions for the rest data by shuffling the sample indices across the entire recording and selecting an equal number of random sample indices per permutation run corresponding to the observed number of burst onsets / respiration peaks. The subsequent analysis steps were identical to the observed data. The shuffled onset / peak times were translated into a phase value relative to the R-peaks before and after each timestamp (see Equation 6) and then the ECG phase coherence, i.e. the mean resultant vector length, for the trial-shuffled reference data was calculated. This procedure was repeated 1000 times for each participant creating a random permutation reference distribution under the null-hypothesis of no trial-wise relationship between burst onset / respiration peak times and cardiac cycles (see Fig. 6C and Supplementary Fig. 1, 4 and 5). As for the observed data, the permutation distributions of ECG phase coherence values were then averaged across participants. The proportion of values in the corresponding reference distribution that exceeded the observed ECG phase coherence value yielded an estimated *p*-value.

## Acknowledgements

We thank Juliane Schubert, Elie Rassi, Tzvetan Popov, Virginie van Wassenhove and Wolfgang Klimesch for very helpful discussions about this work.

## Author contributions

A.W. and N.W. conceived research; A.W. designed research; A.W., M.R. and M.S. performed research; R.A., F.S. and T.H. contributed analytic tools; A.W. and R.A. analyzed data; A.W. wrote the original draft; A.W. wrote the paper.

## Data and materials availability

All data, code and materials will be made publicly available upon acceptance of the paper at the following doi: https://doi.org/10.60817/gcj5-5727

## List of Supplementary Materials

Supplementary Figures S1-S5

**Supplementary Figure S1.**
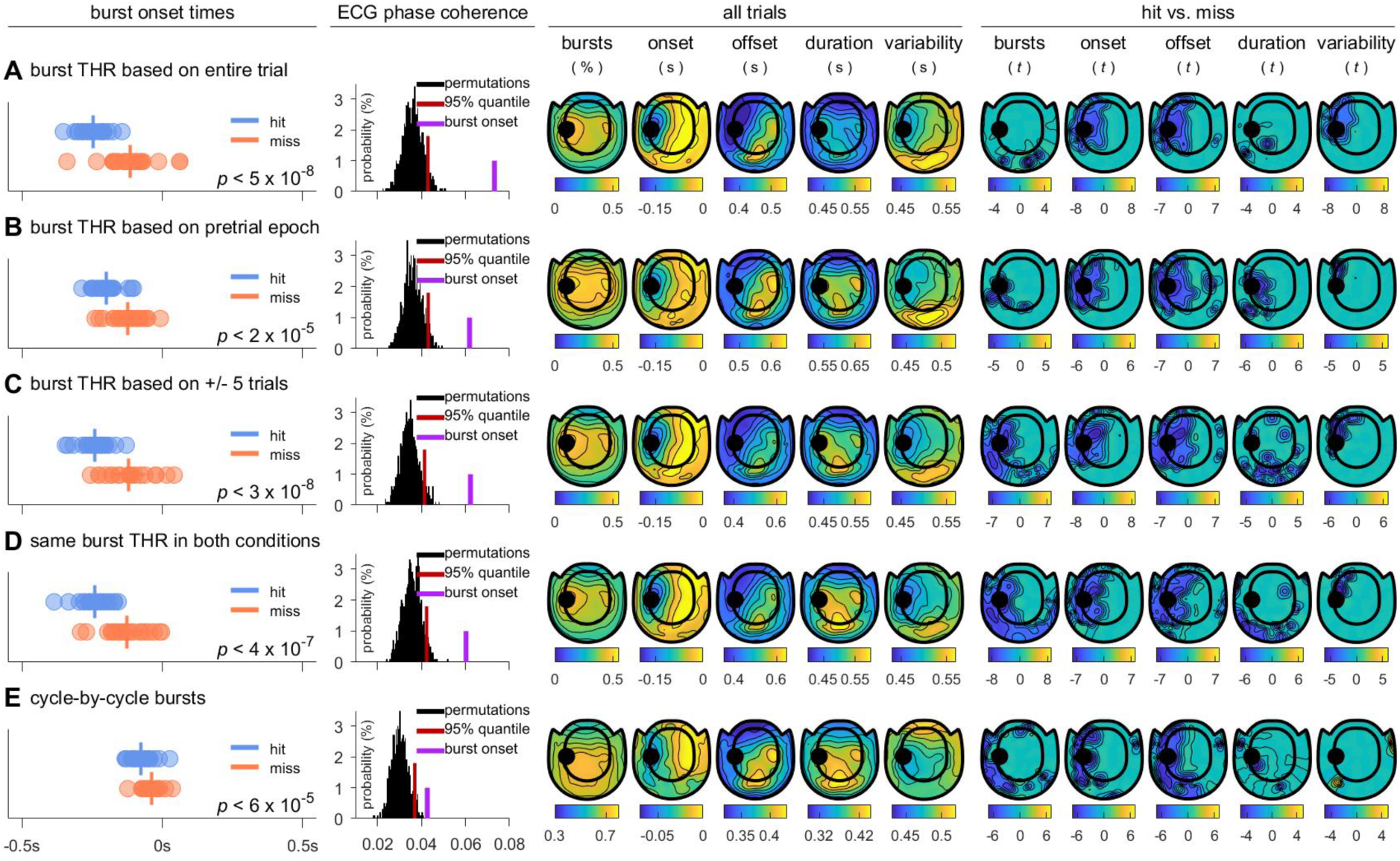
Sensitivity analysis of oscillatory burst detection **A** Burst onset times (left panel) on a representative MEG gradiometer sensor (black dot on sensor-level topographies on the right) per participant and perceptual outcome (blue circles for detected, red circles for missed trials). Vertical lines show the mean onsets. *P*-values denote significance levels for repeated-measures *t*-statistics. Inter-trial ECG phase coherence (middle panel) at burst onset (purple) and for 1000 random permutations. The red vertical line denotes the 95%-quantile of the permutation distribution. Sensor-level topographies (right panel) for all burst occurrences in the interval between -1s to 1s (in percent trials), for the burst onset- and offset times, for the burst durations and for the burst onset variability across trials (all in s), as well as for repeated-measures *t*-statistics between hit- vs. miss-trials for all measures (Bonferroni-corrected). The topographies showing statistics are masked at a corrected threshold of *p* < .05. Panel **A** shows the data for the burst detection method reported in the main text. Panels **B-E** shows the data for different burst detection algorithms using threshold estimates based on the pretrial period (**B**), based on the +/- 5 trials relative to the current trial (**C**), based on the same threshold for both conditions of interest (**D**), and for a waveform-based method (**E**).

**Supplementary Figure S2.**
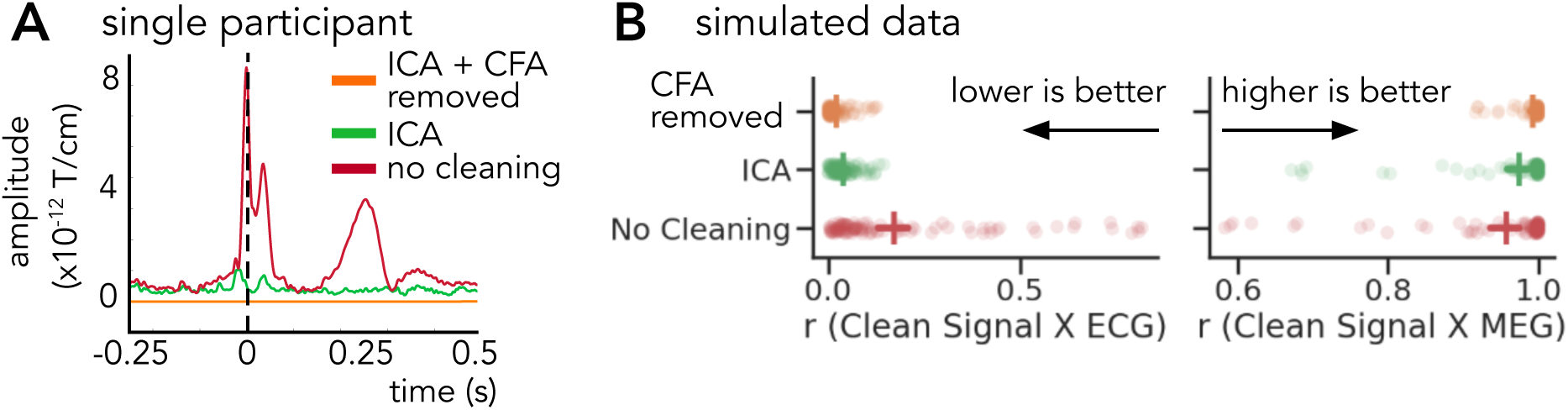
Removal of cardiac field artifacts **A** MEG activity time-locked to R-peaks averaged over all gradiometer sensors without cleaning (red), with ICA-correction (green), and with ICA-correction and removal of the average cardiac field artifact from each trial (orange, used method) for a representative participant. **B** Correlation coefficients between simulated MEG data with an artificially introduced cardiac field artifact with the ECG (left) and with the original MEG (right) for the three different methods.

**Supplementary Figure S3.**
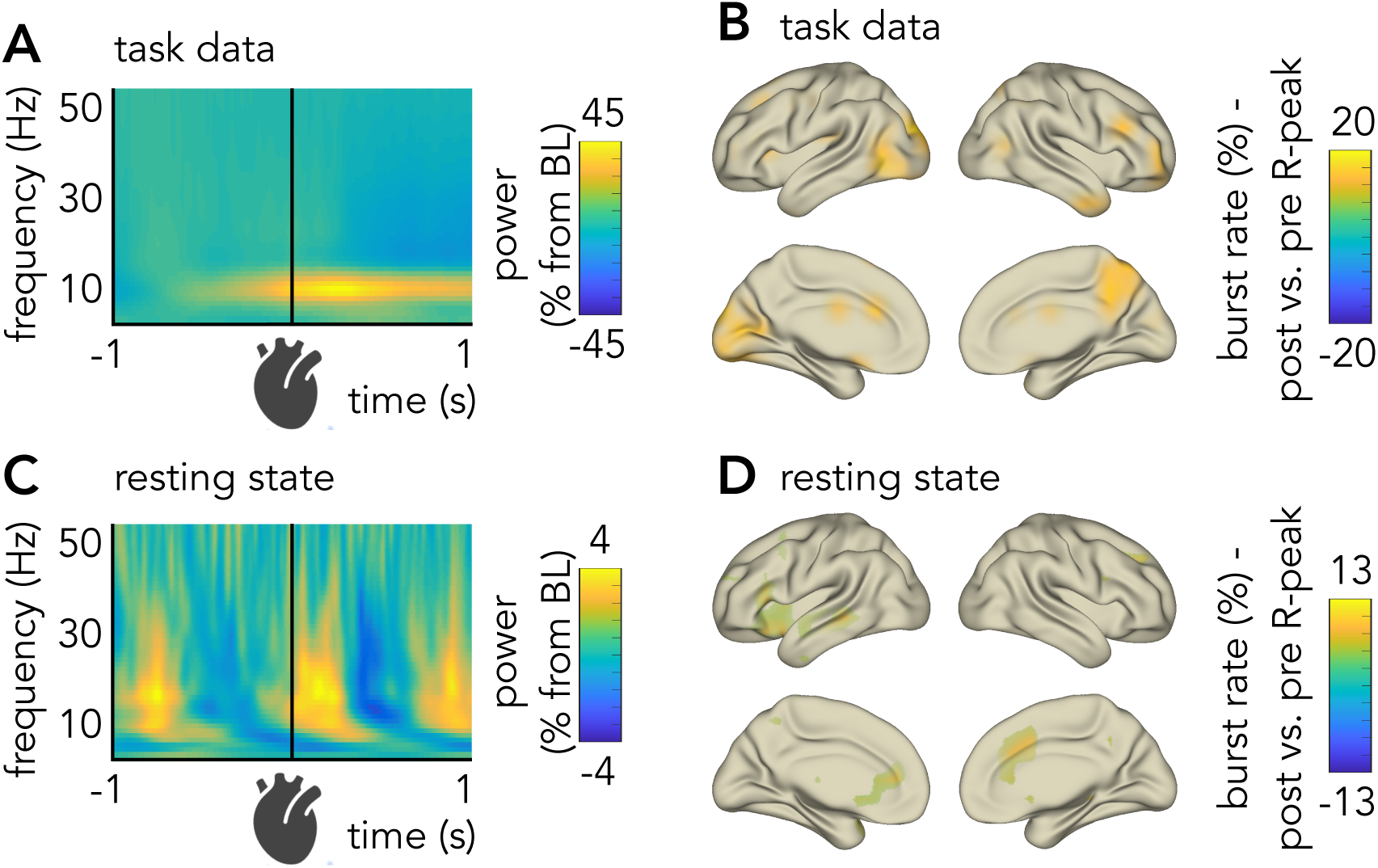
Heartbeat-locked MEG-activity during the task and at rest **A** Time-frequency map for oscillation power vs. baseline time-locked to all R-peaks during the task. **B** Source-level distributions for the burst rate difference between post- vs. pre- R-peaks (pre: -0.3s to -0.1s, post: +0.1s to +0.3s) during the task. **C** and **D** as in **A** and **B** but for resting state data. The upper 10%-voxels are shown.

**Supplementary Figure S4.**
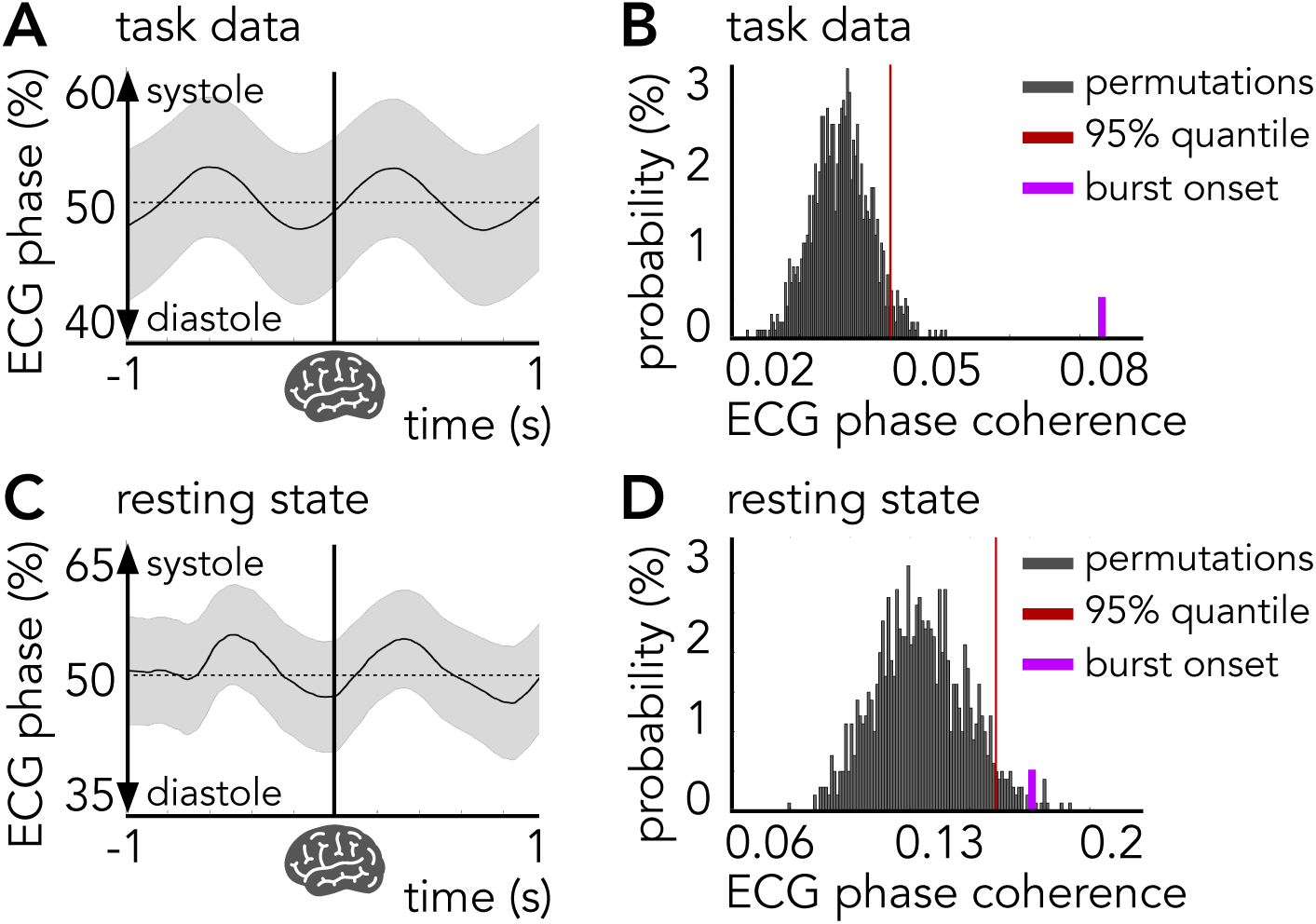
Burst-locked ECG-activity during the task and at rest **A** ECG phase locked to burst onset (straight vertical line) during the task. Increasing values show more systolic-and decreasing values more diastolic activity. Shaded error regions represent SEM. **B** Inter-trial ECG phase coherence during the task at burst onset (purple) and for 1000 random permutations. The red vertical line denotes the 95%-quantile of the permutation distribution. **C** and **D** as in **A** and **B** but for resting state data.

**Supplementary Figure S5.**
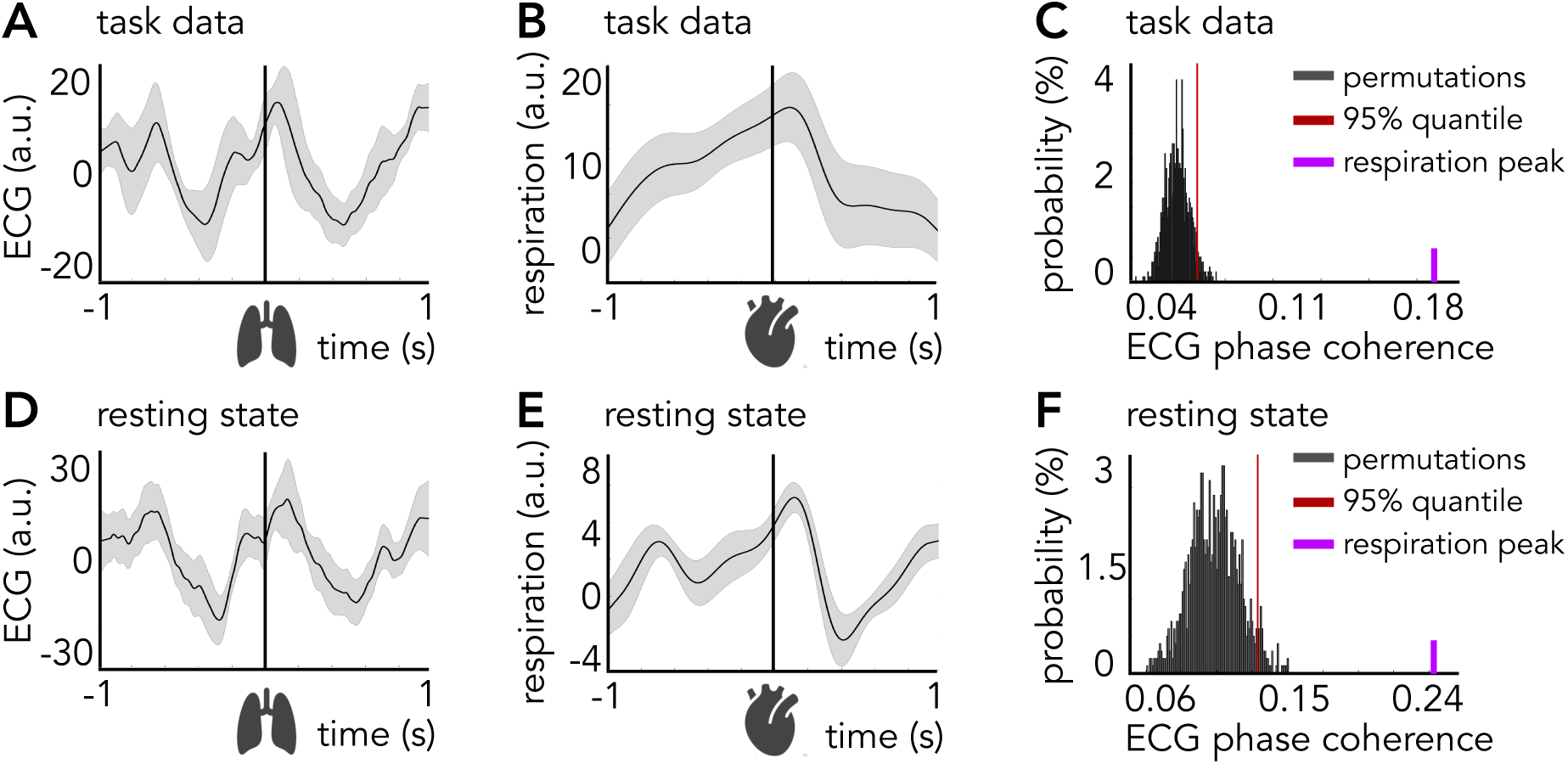
Relationships between ECG- and respiratory activity **A** ECG time-locked to all respiration peaks (straight vertical line) during the task. **B** Respiration signal time-locked to all R-peaks (straight vertical line) during the task. Shaded error regions represent SEM. **C** Inter-trial ECG phase coherence at the respiration peak (purple) and for 1000 random permutations. The red vertical line denotes the 95%-quantile of the permutation distribution. **D-F** as **A-C** but for resting state data.

