## Supplementary Figures for "Brain-body timing gates conscious access across sensory systems"

### List of Supplementary Materials:

Supplementary Figures S1-S5

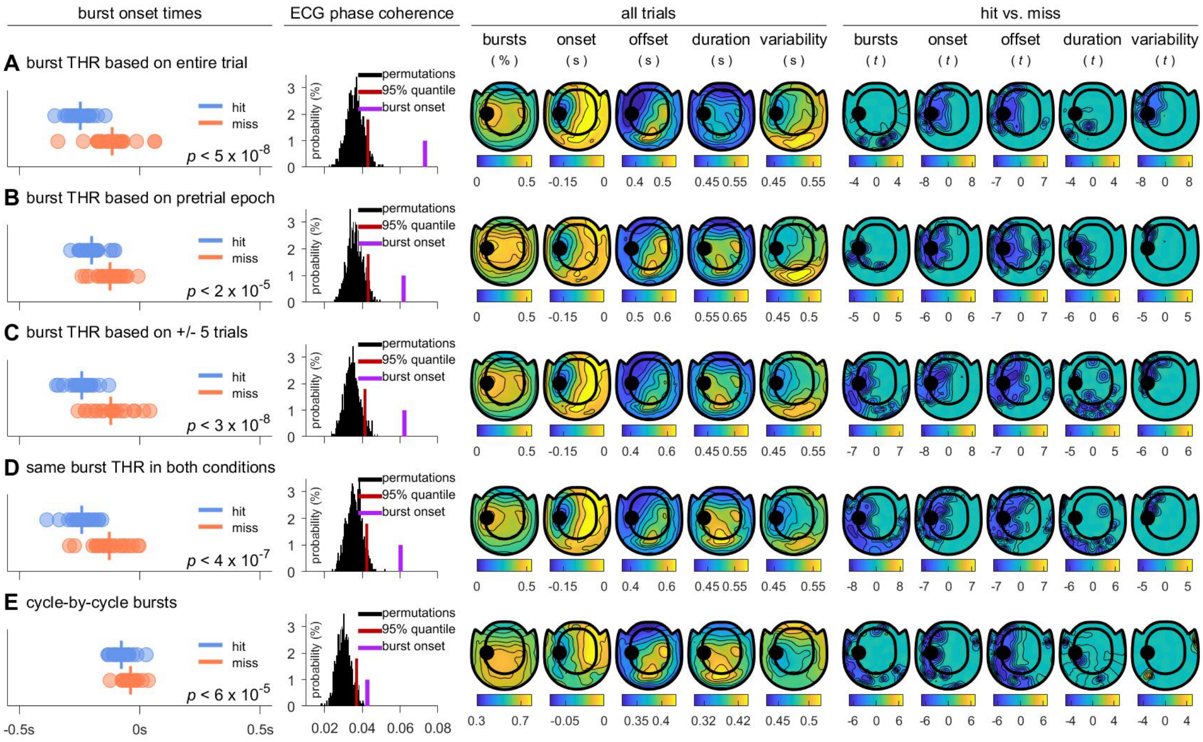


**Supplementary Figure S1.** Sensitivity analysis of oscillatory burst detection

**A** Burst onset times (left panel) on a representative MEG gradiometer sensor (black dot on sensor-level topographies on the right) per participant and perceptual outcome (blue circles for detected, red circles for missed trials). Vertical lines show the mean onsets. *P*-values denote significance levels for repeated-measures *t*-statistics. Inter-trial ECG phase coherence (middle panel) at burst onset (purple) and for 1000 random permutations. The red vertical line denotes the 95%-quantile of the permutation distribution. Sensor-level topographies (right panel) for all burst occurrences in the interval between -1s to 1s (in percent trials), for the burst onset- and offset times, for the burst durations and for the burst onset variability across trials (all in s), as well as for repeated-measures *t*-statistics between hit- vs. miss-trials for all measures (Bonferroni-corrected). The topographies showing statistics are masked at a corrected threshold of $p < .05$. Panel **A** shows the data for the burst detection method reported in the main text. Panels **B-E** shows the data for different burst detection algorithms using threshold estimates based on the pretrial period (**B**), based on the +/- 5 trials relative to the current trial (**C**), based on the same threshold for both conditions of interest (**D**), and for a waveform-based method (**E**).

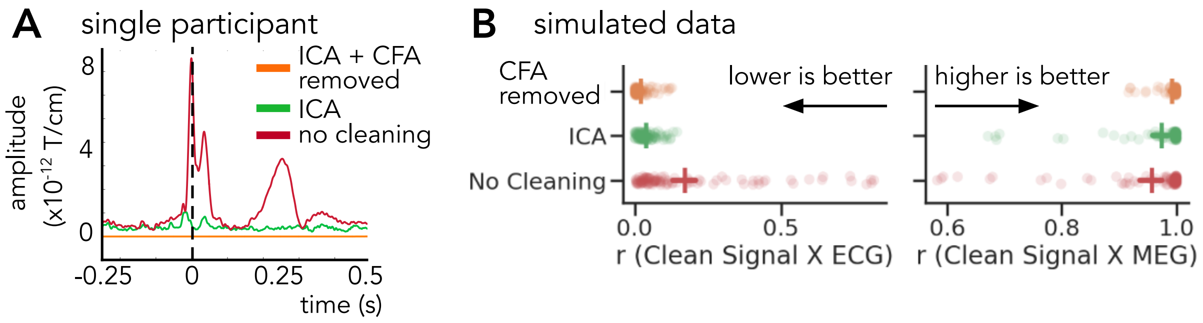


**Supplementary Figure S2.** Removal of cardiac field artifacts
**A** MEG activity time-locked to R-peaks averaged over all gradiometer sensors without cleaning (red), with ICA-correction (green), and with ICA-correction and removal of the average cardiac field artifact from each trial (orange, used method) for a representative participant. **B** Correlation coefficients between simulated MEG data with an artificially introduced cardiac field artifact with the ECG (left) and with the original MEG (right) for the three different methods.

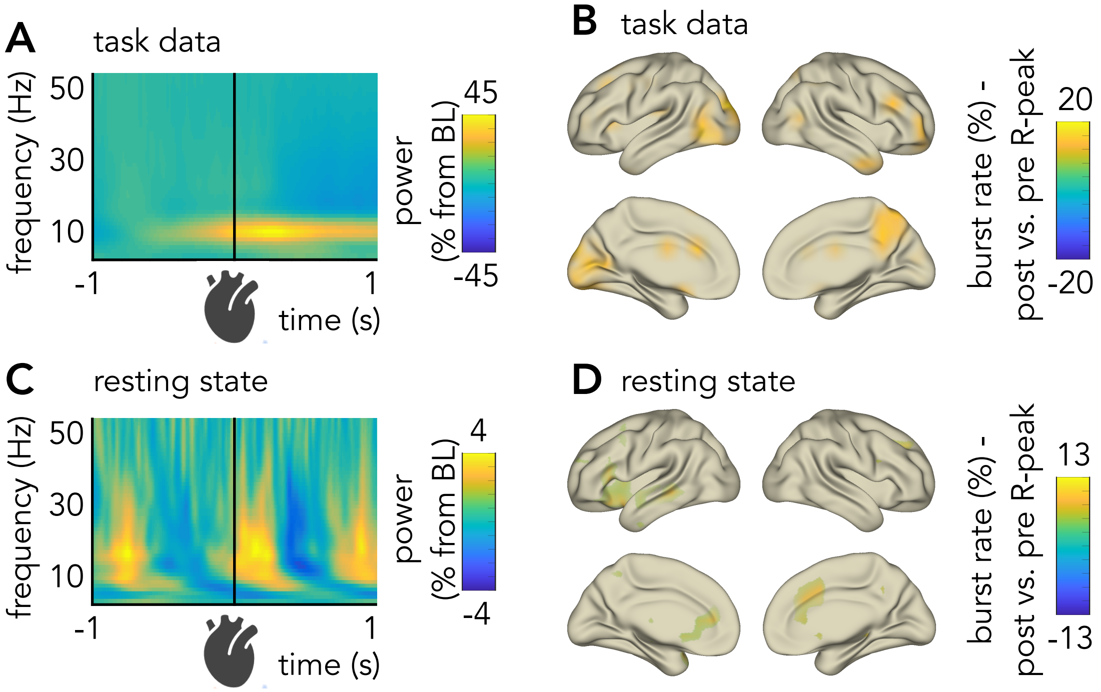


**Supplementary Figure S3.** Heartbeat-locked MEG-activity during the task and at rest
**A** Time-frequency map for oscillation power vs. baseline time-locked to all R-peaks during the task. **B** Source-level distributions for the burst rate difference between post- vs. pre- R-peaks (pre: -0.3s to -0.1s, post: +0.1s to +0.3s) during the task. **C** and **D** as in **A** and **B** but for resting state data. The upper 10%-voxels are shown.

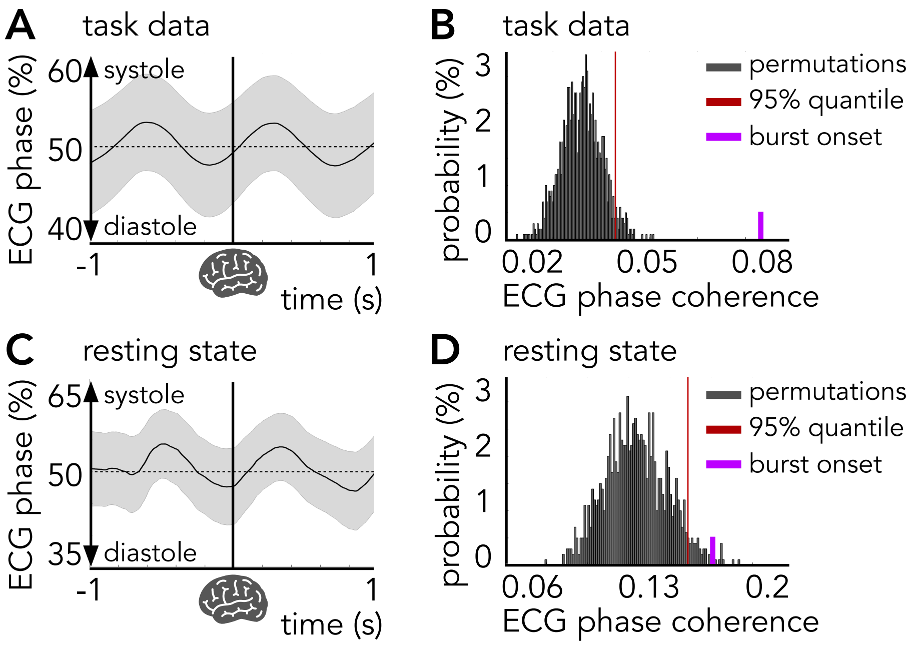


**Supplementary Figure S4.** Burst-locked ECG-activity during the task and at rest
**A** ECG phase locked to burst onset (straight vertical line) during the task. Increasing values show more systolic- and decreasing values more diastolic activity. Shaded error regions represent SEM. **B** Inter-trial ECG phase coherence during the task at burst onset (purple) and for 1000 random permutations. The red vertical line denotes the 95%-quantile of the permutation distribution. **C** and **D** as in **A** and **B** but for resting state data.

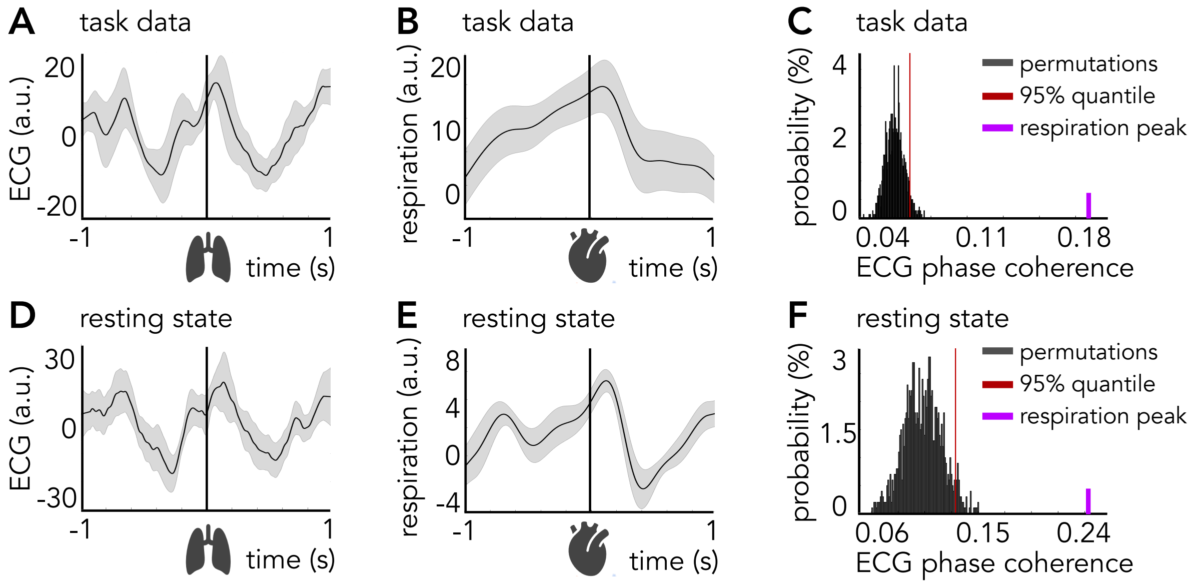


**Supplementary Figure S5.** Relationships between ECG- and respiratory activity
**A** ECG time-locked to all respiration peaks (straight vertical line) during the task. **B** Respiration signal time-locked to all R-peaks (straight vertical line) during the task. Shaded error regions represent SEM. **C** Inter-trial ECG phase coherence at the respiration peak (purple) and for 1000 random permutations. The red vertical line denotes the 95%-quantile of the permutation distribution. **D-F** as **A-C** but for resting state data.
